# Population temporal structure supplements the rate code during sensorimotor transformations

**DOI:** 10.1101/132514

**Authors:** Uday K. Jagadisan, Neeraj J. Gandhi

**Author notes:** **Corresponding author:** Correspondence and requests for more information should be addressed to U.K.J or N.J.G. **Author Contributions:** U.K.J and N.J.G designed the study. U.K.J. performed the experiments, analyzed the data, and performed model simulations. U.K.J and N.J.G wrote the manuscript.

## Abstract

Sensorimotor transformations are mediated by premotor brain networks where individual neurons represent sensory, cognitive, and movement-related information. Such multiplexing poses a conundrum – how does a decoder know precisely when to initiate a movement if its inputs are active at times when a movement is not desired (e.g., in response to sensory stimulation)? Here, we propose a novel hypothesis: movement is triggered not only by an increase in firing rate, but critically by a reliable temporal pattern in the population response. Laminar recordings in the superior colliculus (SC), a midbrain region that plays an essential role in orienting eye movements, indicate that the temporal structure across neurons is a factor governing movement initiation. Specifically, using a measure that captures the fidelity of the population code - here called temporal stability - we show that the temporal structure fluctuates during the visual response but becomes increasingly stable during the movement command, even when the mean population activity is similar between the two epochs. Analyses of pseudo-populations in SC and cortical frontal eye fields (FEF) corroborated this model. We also used spatiotemporally patterned microstimulation to causally test the contribution of population temporal stability to movement initiation and found that stable stimulation patterns were more likely to evoke a movement, even when other features of the patterns such as mean pulse rates and population state subspaces were matched. Finally, a spiking neuron model was able to discriminate between stable and unstable input patterns, providing a putative biophysical mechanism for decoding temporal structure. These findings offer an alternative perspective on the relationship between movement preparation and generation by situating the correlates of movement initiation in the temporal features of activity in shared neural substrates. They also suggest a need to look beyond the instantaneous rate code at the single neuron or population level and consider the effects of short-term population history on neuronal communication and behaviour.

**Summary:** Sensorimotor transformations are mediated by premotor brain networks where individual neurons represent sensory, cognitive, and movement-related information. Such multiplexing poses a conundrum - how does a decoder know precisely when to initiate a movement if its inputs are active at times when a movement is not desired (e.g., in response to sensory stimulation)? Here, we propose a novel hypothesis: movement is triggered not only by an increase in firing rate, but critically by a reliable temporal pattern in the population response. Laminar recordings in the macaque superior colliculus (SC), a midbrain hub of orienting control, and pseudo-population analyses in SC and cortical frontal eye fields (FEF) corroborated this hypothesis. Importantly, we used spatiotemporally patterned microstimulation to causally verify the importance of temporal structure and demonstrate its role in gating movement initiation. We also offer a spiking neuron model with dendritic integration as a putative mechanism to decode this temporal information. These findings offer new insights into the long-standing debate on movement generation and highlight the importance of short-term population history in neuronal communication and behavior.

## INTRODUCTION

In order to successfully interact with the environment, the brain must funnel down the sensory inputs it receives to specific movements at specific times. Such sensory-to-motor transformations are critically mediated by premotor brain networks where evolving activities in individual neurons represent sensory, cognitive, and movement-related information (*1−3*). For example, in brain regions involved in the control of gaze, including the superior colliculus (SC) and the frontal eye fields (FEF), a population of visual neurons processes the stimulus, while another group of motor or movement neurons discharges a volley of spikes to produce the saccade to the location of the stimulus. So-called visuomovement neurons, which constitute the majority, burst for both stimulus presentation and saccade generation (*1*). In reality, visuomovement and motor neurons span a continuum that exhibits different amounts of visual and movement-related activities. Such neurons from both SC and FEF project to the saccade generation circuitry in the brainstem (*4, 5*). How the activity patterns of visuomovement and motor neuron populations collectively drive saccade initiation remains an unresolved question in motor systems neuroscience.

Canonically, accumulation to threshold mechanisms have been thought to underpin saccade initiation (*6, 7*). The accumulator model posits that population activity in SC and FEF increases stochastically toward a constant threshold activation level, leading to the generation of a high-frequency burst and execution of a saccade. While support for the constant threshold criterion has been reported for FEF motor neurons (*6*), visuomovement neurons in FEF as well as visuomovement and motor cells in SC do not seem to obey the criterion (*8*). Moreover, in many visuomovement neurons, the threshold activation level estimated from the premotor burst is also crossed by the sensory response, yet a movement is not triggered early. It has also been suggested that the threshold criterion applies to the population response, not necessarily to every neuron in the ensemble (*9*). Even then, it remains possible that the population activity could exceed the threshold level during the sensory response. Thus, the accumulator model has limitations.

Variability of neural population activity has also been proposed as a potential mechanism of movement initiation, notably in the skeletomotor system. For simultaneously recorded neural populations, e.g., in premotor or motor cortex, the ensemble response for each trial can be represented as a neural trajectory in multi-dimensional population activity space. These trajectories have been shown to traverse through “optimal subspaces”, defined as regions of reduced variability computed across trials (*10*). In the delayed reach task, for example, such optimal subspaces can be defined for stimulus onset, motor preparation and movement execution stages (*11*). A closely related model posits that muscles are recruited and a movement is initiated when neural activity traverses an optimal or “motor-potent” region of the population state space (*12, 13*) and is inhibited otherwise, e.g., during movement preparation (*13,14*). A major strength of these approaches is that the firing rate need not reach a threshold to trigger the movement (but see (*15*)), yet this framework also has limitations. Since variability measures are computed by combining information across trials, they are not available to be used by downstream structures on individual trials. More importantly, neural variability models do not provide direct mechanistic insights into how neurons decoding their inputs map the presence of a trajectory in a particular subspace into a specific function, nor have they been validated using causal manipulations.

In this study, we propose a novel mechanism for movement generation based on the temporal structure of neural population activity. Our central hypothesis is that saccade initiation occurs when high firing rate (burst) is coupled with stable temporal structure in population activity. We specifically propose that the visual burst is unstable and therefore cannot trigger a saccade, while the premotor burst is stable and does initiate the movement when activity crosses threshold. We demonstrate the presence of this code using a combination of laminar population recordings in the SC and pseudo-population analyses in SC and FEF, suggesting that temporal stability features permeate through the entire visuomotor neuraxis, encompassing both cortex and subcortex (*1*). We also show that the code is robust across tasks and sub-regions of SC. Crucially, we provide causal validation of this mechanism by delivering temporally structured or unstructured microstimulation patterns into the SC with a linear electrode array and comparing their movement-inducing efficacy. Finally, we show how population temporal structure could be decoded by a spiking neuron model with dendritic integration. We foresee this computation as a pervasive feature of neuronal communication, and we apply it specifically to understand the mechanisms of saccade initiation.

## RESULTS

### Joint representation of visual stimulus and premotor processing in visuomovement neurons

The aforementioned, dual nature of visuomovement neurons is best illustrated by examining their activity in the delayed response paradigm (left panel in Figure 1A), which requires the subject to withhold a saccade to a stimulus in the visual periphery until after the disappearance of a central fixation cue. We recorded population activity from the SC using laminar microelectrode arrays (right panel in Figure 1A) in two monkeys (*Macaca mulatta*). Figures 1B shows responses on individual channels (top row) and population activity averaged across channels (bottom row) aligned on target (left) and saccade (right) onsets for three example trials in one session. The population exhibits a high-frequency visual burst following target onset and a subsequent premotor burst prior to a saccade. The peak magnitude of the visual burst is lower than that of the premotor burst on some trials (light traces in Figure 1B), but it is not uncommon for the peak visual response to match or exceed the premotor activity (medium and dark traces in Figure 1B), especially when accounting for the 10-20 ms efferent delay from neural initiation to movement onset (*6, 16, 17*) (vertical dashed line in bottom panel of Figure 1B, also see Figure 1G), consistent with previous reports that tectoreticular neurons involved in saccade generation show strong visual and premotor transients (*4*). Yet, on these trials, the visual burst does not trigger a movement, an observation that casts doubt on thresholding (*6, 8*) as a singular mechanism governing movement initiation.

**Figure 1.**
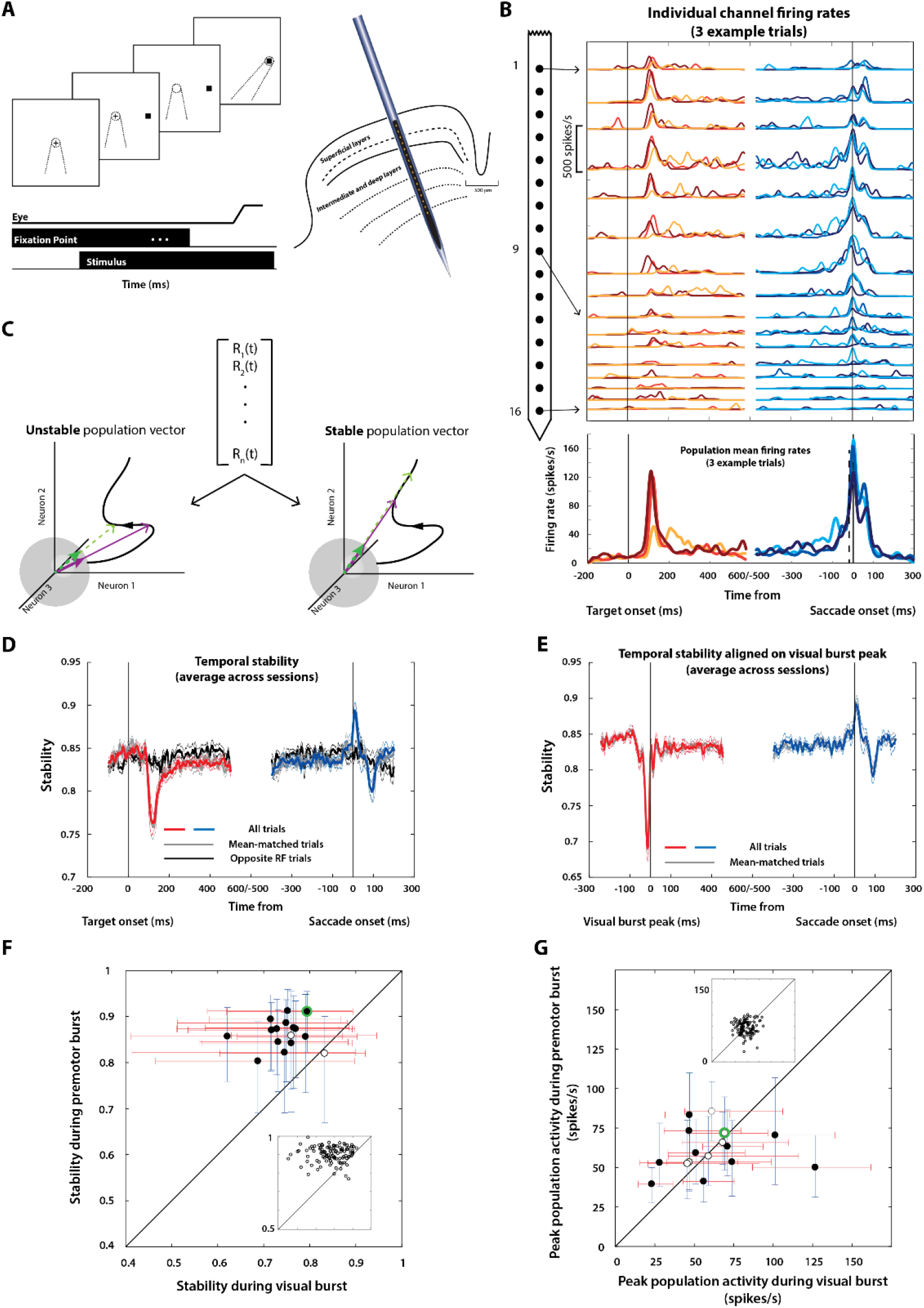
Temporal structure of population activity during sensorimotor multiplexing. **A**. Left - Sequence of events in the delayed saccade task. The top row shows a typical display sequence, and the bottom rows show the timeline. The fixation target is shown as a plus symbol for illustration purposes. Dotted lines depict line of sight of the animal. Right - Linear electrode arrays with 16-24 contacts were used to penetrate the SC normal to the surface. Thus, all neurons encountered along the dorso-ventral axis of the SC prefer similar optimal vectors. **B**. Top: Example recording session showing activity from 16 channels aligned on target (left column) and saccade (right column) onsets for 3 different trials (traces with different color saturations). Bottom: Population activity averaged across the 16 channels for the 3 example trials. Note the considerable variation in the amplitude of the population visual burst across trials. **C**. Schematic depicting computation of temporal stability. The population vector (center) was used to construct a single-trial neural trajectory, which was then normalized at each instance to represent a unit length population vector traversing along a hypersphere. The left and right panels highlight unstable and stable parts of a schematic trajectory. Note that in both cases, the two population vectors (thin arrows in green and purple) are meant to be separated by a similar length of time. In the unstable case, the normalized vectors (thick arrows) move around on the surface of the hypersphere (green and purple vectors are spatially separate), whereas in the stable case, they stay pointed roughly in the same direction (the two vectors overlap in space). **D**. Temporal stability is computed as a dot product between two normalized population vectors separated by 2*τ* = 20 *ms* aligned on target (left) and saccade (right) onsets. The stability of population activity, averaged across sessions, drastically decreases during the visual burst (red traces, mean +/- s.e.m.; p < 0.01 for Wilcoxon signed-rank test with respect to baseline) and increases before and during movement onset (blue traces; p < 0.01 for Wilcoxon signed-rank test with respect to baseline). The gray trace is the stability computed only for those trials where the peak visual burst matched or exceeded activity during the premotor burst at saccade initiation (after accounting for an efferent delay of 15 ms). It is effectively identical to the red and blue waveforms. The black trace shows population stability on trials in which the target and saccade were directed to the opposite hemifield, far away from the response field (RF) of these neurons. **E**. Temporal stability aligned to peak visual burst. The peak time was computed on the average population response of each trial, and the peak-aligned trace was averaged across trials and sessions. Note the sharper dip in the stability of the visual response when aligned to the peak. Like in D, the gray trace shows peak-aligned stability for trials in which the peak visual activity matched or exceeded premotor activity at saccade initiation. The gray and colored traces are nearly identical. **F**. Scatter plot of temporal stability in a 20 ms window preceding the peak of visual and premotor bursts for individual sessions. Each point is the median of the stability values computed on individual trials for one session (error bars are +/- 95% confidence intervals). Filled circles indicate significant difference between the visual and premotor stability (p < 0.01, Wilcoxon signed-rank test). The inset shows the scatter plot for individual trials for an example session (encircled in green). Thus, temporal stability of visual and premotor epochs on individual trials also showed the same effect. **G**. Similar to F but for the peak population firing rate during the visual and premotor bursts. The inset is for the same example session as in F. For F and G, the premotor burst is truncated at 15 ms before saccade onset, the putative efferent delay between SC and the extraocular muscles.

We also tested whether saccade initiation could be controlled solely by “motor” neurons, thought to be active only during saccades and not during the visual epoch. This hypothesis requires motor neurons to exclusively have a high motor potential, i.e., be most correlated with saccade kinematics (*18*). While motor neurons indeed have a high motor potential (Supplementary Fig. 1A and B), we found that neurons with high visual activity also were among the strongest drivers of the saccade (Supplementary Fig. 1C and D), disputing the categorical motor neuron hypothesis. Given that visuomovement and motor neurons project directly to the brainstem saccade burst generator that initiates and guides saccadic gaze shifts (*4, 5, 19*), we asked how downstream structures are able to differentiate between the two bursts.

### Population temporal structure is scrambled during the visual response and stable during the premotor response

We reasoned that if static features of population activity such as the mean firing rate are insufficient to discriminate between visual and premotor bursts, the answer likely resides in the spatiotemporal structure of activity during the two bursts. Indeed, a growing body of work has revealed that precise coordination in the timing of input spikes is more efficient at driving downstream cortical and subcortical neurons (*20-22*). Specifically, we hypothesized that a critical criterion for a decoder to generate a movement should be a high level of certainty in the instructed movement and its metrics. Since the exact decoding scheme is unknown, we considered whether certainty is provided by consistency in the population pattern over the course of a burst, under any generalized weighted pooling scheme. To quantify this consistency, we first constructed a population firing rate trajectory as a function of time. Next, we normalized the population vector at each time point, constraining it to a unit hypersphere in state space (Figure 1C; see Methods for details). This step factors out global changes in firing across the population and focuses on the relative activity pattern. We then computed the dot product between two of these unit vectors separated in time (parametrized by *τ*) - we call this measure the temporal stability of the population code.

Figure 1D shows the evolution of temporal stability averaged across all sessions (n = 280 channels/16 sessions, mean +/- s.e.m., colored traces; also see Supplementary Fig. 2A, top row). The stability of the population pattern decreased relative to baseline during the visual burst and increased during the premotor burst. Moreover, this property was preserved when considering only the subset of trials where the peak visual response matched or exceeded the premotor activity 15 ms before saccade onset (gray trace in Figure 1D) and was significantly different from the trend in the null condition when the saccade was directed to a stimulus in the opposite hemifield (black trace in Figure 1D). Scrambling the population code by either shuffling the activations of individual neurons at each time point or shuffling across time points lowered the stability profile for the entire trial significantly (Supplementary Fig. 2A, bottom row). The anti-phase relationship of stability between the visual and premotor bursts was also present across a range of separation times between the two population vectors used for the dot product (Supplementary Fig. 2B). We also computed temporal stability by realigning activity on the peak population visual response, to discount any effect of visual response onset latency differences across the population and found similar (if not stronger) effects (Figure 1E, also see Supplementary Fig. 2C). Finally, temporal stability afforded more robust discrimination between the visual and premotor bursts even on individual trials and sessions (Figure 1F), compared to the population firing rate (Figure 1G). These results were unaffected by spike sorting (Supplementary Fig. 3, also see Methods), in line with recent evidence that spike sorting has a negligible effect on population analyses (*23*). Thus, population temporal stability seems to impose a constraint that prevents movement initiation at an undesirable time (visual epoch), allowing the animal to successfully perform the task at hand. Once cued to execute a gaze shift, the activity in the same population rises in a stable manner allowing saccade initiation.

### The distinction between visual and premotor temporal structure is robust across brain regions and tasks

Next, we explored the robustness of the temporal stability hypothesis. We tested whether the properties of stability observed in other populations of neurons or conditions were consistent with its predicted role in movement initiation. For this part of the study we used data obtained from single-electrode recordings in SC and the frontal eye fields (FEF), the latter of which plays a major role in the cortical control of saccade initiation (*6, 24*). First, we constructed pseudo-trials by random sampling of trials from individual neurons/single-unit recording sessions (see Methods for details) and used this pseudo-population to construct surrogate neural trajectories (Figure 2A, top row, also see Methods and Supplementary Fig. 4). Since SC and FEF both project to saccade-generating brainstem structures (*25*), we also analyzed a combined SC+FEF pseudo-population. The middle row of Figure 2A shows the average normalized pseudo-population activity of 60 SC neurons, 30 FEF neurons, and the combined 90-neuron population, reaffirming the visuomovement pattern of activity discussed earlier. Importantly, the temporal structure profiles (Figure 2A, bottom row) were remarkably similar to those observed with simultaneous population recordings – the dramatic reduction in the stability during the visual burst and an increase in stability during the premotor burst – in both SC and FEF individually (transparent and opaque colored traces in Figure 2A, bottom row), as well as in the combined population (black traces in Figure 2A, bottom row).

**Figure 2.**
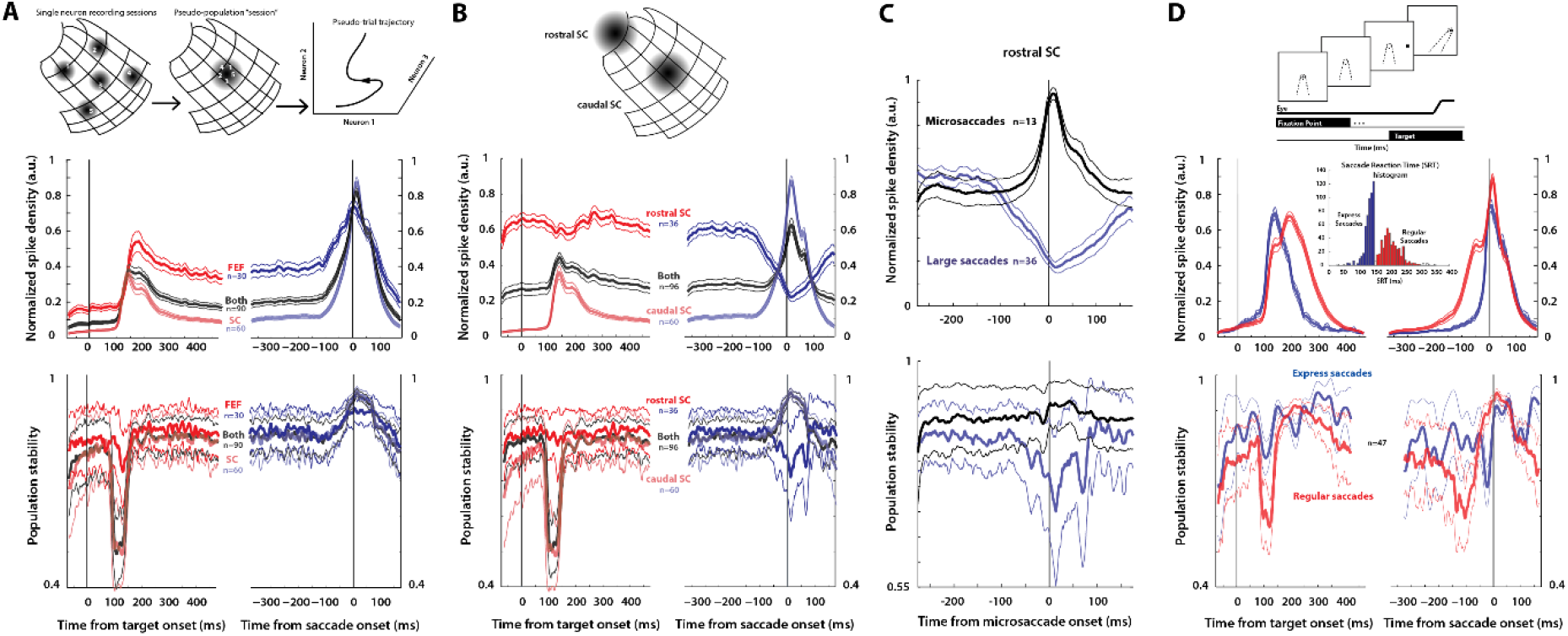
Temporal stability in SC and FEF pseudo-populations. **A**. Top row – Pseudo–population schematic. We combined single–neuron recording trials from different active populations on the SC map (left) into one pseudopopulation (middle) that would be active for an average saccade, enabling the construction of a pseudo–population trajectory. A similar construction (not depicted) was performed for FEF. Middle row – Mean (+/– s.e.m.) normalized pseudo–population activity of visuomovement and motor neurons in SC (light red and blue traces) and FEF (dark red and blue traces) during the delayed saccade task. The black traces show the combined SC+FEF population putatively jointly projecting to the brainstem. The left and right panels show data aligned on target and saccade onsets, respectively. Bottom row – Temporal stability of the SC, FEF, and combined SC+FEF populations (colors as in the middle row). Pseudo–population stability shows a drastic reduction at the time of the visual burst and an increase prior to the onset of movement, consistent with the simultaneously recorded laminar population data in the previous figure. **B**. Top row – Schematic depicting active population regions in rostral and caudal SC. Middle row – Mean normalized pseudo–population activity in rostral SC neurons during the delayed saccade task (dark red and blue traces), overlaid with caudal SC activity (light red and blue traces as in panel A) and the combined rostral+caudal SC pseudo–population (black traces). Bottom row–Temporal stability of the rostral, caudal, and combined rostral+caudal pseudo–populations. Note that activity suppression in the rostral SC for large saccades is associated with a concomitant reduction in temporal stability. **C**. Population activity of rostral SC neurons in stable during microsaccades. Top row – Normalized population activity of microsaccade–related neurons in rostral SC (black trace). Note the strong burst compared to the suppression for large saccades in the same population (blue trace). Bottom row – Temporal stability of the rostral SC population for microsaccades and large saccades. Neurons which burst for microsaccades remain stable during these fixational eye movements, consistent with the hypothesis that both and increase in firing rate and high stability is required for movement generation. **D**. Top row – Sequence of events in the Gap task, designed to elicit express saccades. Middle row – Mean normalized pseudopopulation activity on regular (red traces) and express (blue traces) saccade trials, as denoted in the inset of reaction time distribution. Bottom row – Population temporal stability decreases during the visual burst on regular saccade trials (red traces) like in the delayed saccade task but remains relatively high on express saccade trials in the same time period.

We then used pseudo-population analyses to test whether the temporal stability framework was obeyed by SC neurons beyond those that are involved in the generation of large or non-fixational saccades. Neurons in the rostral SC are tonically active during fixation and reduce their activity during larger movements (*26*) (Figure 2B, middle row). Importantly, they also project to the saccade burst generator (*27*). Assuming population stability modulates the input drive to downstream structures, we hypothesized that the temporal structure in rostral SC neurons must decrease during large saccades but remain elevated during fixation, even when a visual burst occurs in other parts of SC and FEF. In other words, we expected the evolution of temporal stability across rostral SC neurons to be the inverse of what occurs in caudal SC - Figure 2B (bottom row) confirms this prediction, both when rostral SC is considered separately, as well as in the combined SC population. In addition to the antiphase relationship with caudal SC during large saccades, rostral SC is also known to play a causal role in the generation of microsaccades (*28*) during fixation. Therefore, we asked whether the rostral SC population exhibits stable temporal structure during the burst that produces microsaccades. This was indeed the case, where the same rostral SC population that showed a reduction in temporal stability during large saccades showed an increase during microsaccades (Figure 2C). Thus, the population activity of neurons in rostral SC, including its temporal structure, complements the pattern in other parts of SC in both suppressing and initiating movements.

We also examined whether population temporal structure was related to the propensity to trigger saccades off the visual burst. In a modified task paradigm called the gap task (top row, Figure 2D), “express saccades” occur under certain conditions, and are reflected in the initial peak of a bimodal reaction time distribution (inset in middle row, Figure 2D). These low-latency saccades are thought to be triggered directly by the initial visual burst in SC (middle row, Figure 2D). If temporal structure of SC activity is related to the saccade generation process, then temporal stability should not decrease at all or as much during the visual burst when express saccades are triggered. The bottom row of Figure 2D shows that this was indeed the case – the temporal stability in the visual epoch is higher for express saccades (it drops slightly but quickly recovers) compared to the notable trough for regular saccades. In sum, temporal stability seems robustly correlated with saccade generation across brain regions (rostral SC, caudal SC, and FEF), tasks (delayed, regular, and express saccades), experimental techniques (laminar and single electrode recordings), and analytical approaches (spike sorted, unsorted, and pseudo-population analyses).

### Microstimulation patterns with stable temporal structure are more likely to evoke movements

Thus far, we have shown that population temporal structure is a candidate measure that can be used to discriminate between visual and premotor bursts, using purely correlational analyses. To test whether this measure is actively used by the brain in a causal manner, we performed multi-site patterned microstimulation in SC. We first identified individual contacts of the laminar probe that evoked low-latency saccades with suprathreshold stimulation, to verify their placement in the intermediate layers of SC. We then designed temporally stable and unstable stimulation patterns restricted to these contacts, limiting stimulation parameters for individual sites to the sub-threshold regime and verifying that the overall stimulation was near-threshold (for details, see Methods). Stable patterns were created with a linearly decreasing inter-pulse interval (IPI) sequence for one contact (*29*) (to simulate a burst) and scaling the IPIs for other contacts by a uniformly spaced factor. For each stable pattern, we also created a *paired* unstable pattern by jittering the pulse times within a window and shuffling spikes between contacts, preserving both the pulse count for each site and the average pulse rate across the “population”.

An example pair of stable-unstable pulse trains is shown in Figure 3A (top row). The pulse rates determined from these trains (second row) illustrate the scaled rates for the stable pattern and the fluctuating rates for the unstable pattern for individual sites, despite the comparable population rates on individual trials (thick traces; see Supplementary Fig. 5B,C for more examples). Note that the assignment of high and low rates to different contacts was randomized across trials, resulting in similar trial-averaged pulse rates for any given channel, for either type of pattern (third row). The temporal stability for stable and unstable patterns for all trials in the example session (bottom row) confirm the impression provided by the pulse trains and rates, i.e., the scaled patterns are highly stable compared to the relative instability of the jittered patterns.

**Figure 3.**
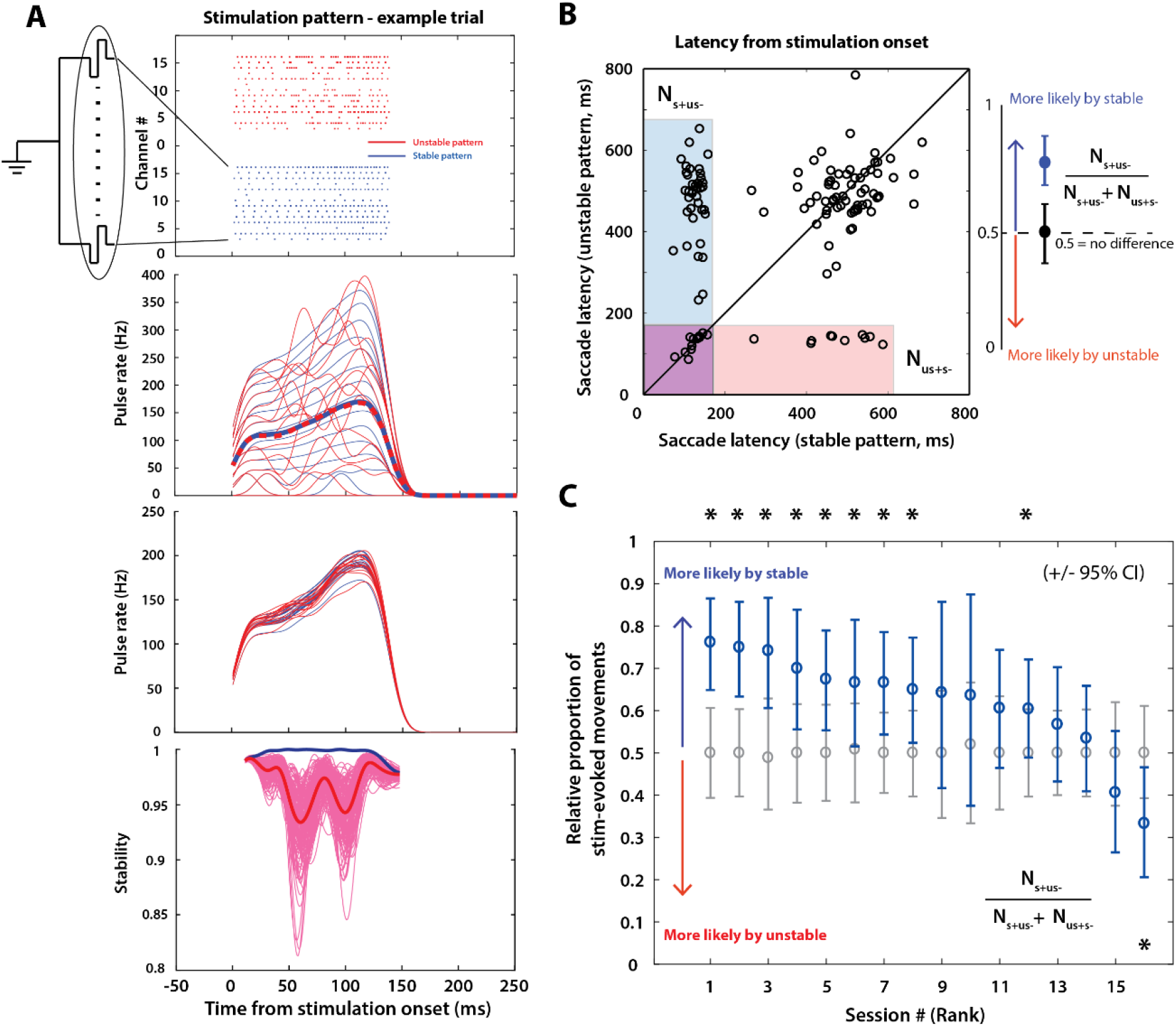
Patterned microstimulation supports temporal stability as a model of movement initiation. **A**. Example pair of stable-unstable stimulation patterns. Top row - Pulse trains for 14 of 16 channels that were stimulation–viable with suprathreshold parameters for this session. Red and blue trains indicate an example pair of unstable and stable patterns, respectively. Second row–Pulse rates for the 14 channels in the stable-unstable pulse patterns, with the population rates averaged across channels overlaid as thick, dashed traces. Note that the rates in the unstable pattern are highly fluctuating despite the matched population average. Third row -- Average pulse rates for each channel across all trials for the stable (blue) and unstable (red) patterns used in this session. The clustering of trial-averaged rates in a narrow band is the intended result after randomization of pulse rate assignment to individual channels on different trials. Bottom row–Temporal stability of the stable and unstable patterns (blue and pink, respectively) for all trials in the example session. Across-trials averages are shown as thick traces. **B**. Scatter plot of saccade latencies (relative to stimulation onset) for an example session. Each point reflects the outcome of stimulation with stable and unstable pulse patterns constituting each pair. The points in shaded regions are stimulation-evoked for at least one condition. N_s+us−_, points in the light blue shaded box, counts trial pairs for which a stable stimulation pattern evoked a saccade but its unstable counterpart did not. N_us+s−_, points in the light red shaded box, denotes the trial pairs for which an unstable stimulation pattern evoked a saccade but its stable counterpart did not. Points in the purple box show subset of trials in which both stable and unstable pairs evoked a saccade. Latency values are comparable for near-threshold stimulation para meters (*53, 54*). Points in the unshaded (white) region refer to trials in which neither stable nor unstable pattern of the pair evoked a saccade. For these pairs, the trial was assigned the latency of the saccade (relative to stimulation onset) directed to a target presented after the gap period. The graphic on the right shows the calculation of relative likelihood of the stable pattern evoking a stimulation-evoked movement for a given session. **C**. Relative proportions of the stable pattern evoking a movement for each session (blue points, error bars represent 95% Cl of the bootstrapped distribution). The sessions are sorted in descending order of proportion for viewing clarity. Note that this sorting order is used in all subsequent figures depicting individual sessions (Figure 4D and Supplementary Fig. 6B). Gray points and error bars are computed from a surrogate dataset in which stable/unstable trial identities are completely shuffled (mean +/- 95% Cl). Asterisks denote a significant effect for that session (p < 0.01 on the permutation test).

We delivered these stimulation patterns during the “gap period” in a gap saccade task (Supplementary Fig. 5A). Figure 3B shows a scatter plot of the stimulation-aligned saccade latencies observed using stable versus unstable stimulation patterns for all pairs from one session. For the majority of stable-unstable trial pairs, a saccade was evoked with the stable but not the unstable pattern, henceforth referred to as S+US-trial pairs. These are the points in the blue shaded window spanning the stimulation duration in Figure 3B. Points in the unshaded region refer to trials in which neither stable or unstable pair evoked a saccade, and these trials were assigned the latency relative to stimulation onset of the saccade directed to a target presented after the gap period. The relative likelihood of evoking a movement with the stable stimulation pattern only, as computed in the graphic on the right, is shown as a function of session number in Figures 3C (blue points). For most sessions, the observed relative likelihood values were significantly higher (i.e., biased towards the stable pattern; p < 0.01, permutation tests) than the null distribution (gray points) obtained by shuffling trials with randomly assigned stable-unstable identities, reinforcing the observation that the stable pattern was more likely to evoke a movement compared to a *state-matched* unstable pattern. We also analyzed the movement vectors evoked by stable and unstable patterns – both sets of saccades, when they occurred, were similar to each other in both amplitude and direction and to the site-specific vectors evoked by constant frequency multi-channel stimulation (Supplementary Fig. 5D and 5E; p > 0.01 for most sessions, Wilcoxon rank-sum tests; for details, see Methods).

Additionally, we explored whether differences in pulse rates on subsets of trials could explain the across-sessions range of movement likelihood effects seen in Figure 3C (Supplementary Fig. 6). Sessions in which the stable pattern was more likely to evoke movements were also the ones in which the S+US-trials (Supplementary Fig. 6A) were closest to the intended uniform sampling of pulse rates across channels, as reflected in the low relative dispersion index for those sessions (Supplementary Fig. 6B and 6C). Sessions with no significant (or negative) effect on movement likelihood were the ones which had a high across-channel pulse rate sampling asymmetry, potentially allowing extremes in pulse rates to mask any underlying effect of temporal structure (Supplementary Fig. 6B and 6C; also see Methods).

### Linear decoders using temporal information outperform static models of movement generation

Having demonstrated the role of population temporal structure in saccade initiation using both correlative and causal approaches, we sought to disambiguate the stability framework from extant models of movement initiation. We earlier argued for the implausibility of threshold-based gating based on the fact that the population activity during the visual burst can sometimes exceed premotor activity (Figure 1B & 1G). Could other population activity-based mechanisms, such as weighted linear readout models that seem to govern movement generation in the skeletomotor system (e.g., the optimal or movement-potent subspace frameworks) (*12-14*), play a role here? To test this, we first used factor analysis (FA) to visualize the low-dimensional neural states (*30*) in the visual and premotor bursts (see Methods). An example 3-dimensional FA projection of the two bursts is shown in Figure 4A – the two sets of states are clearly separable, likely an effect of the subtle yet distinct trends in visual versus premotor activity levels along the dorso-ventral extent of the SC (e.g., Figure 1B) (*31*). Indeed, a linear discriminant analysis (LDA) classifier was able to easily discriminate between the visual and premotor states for most sessions (histogram inset in Figure 4A). The properties of neural activity in SC therefore seem to be consistent with a static state space code as well.

**Figure 4.**
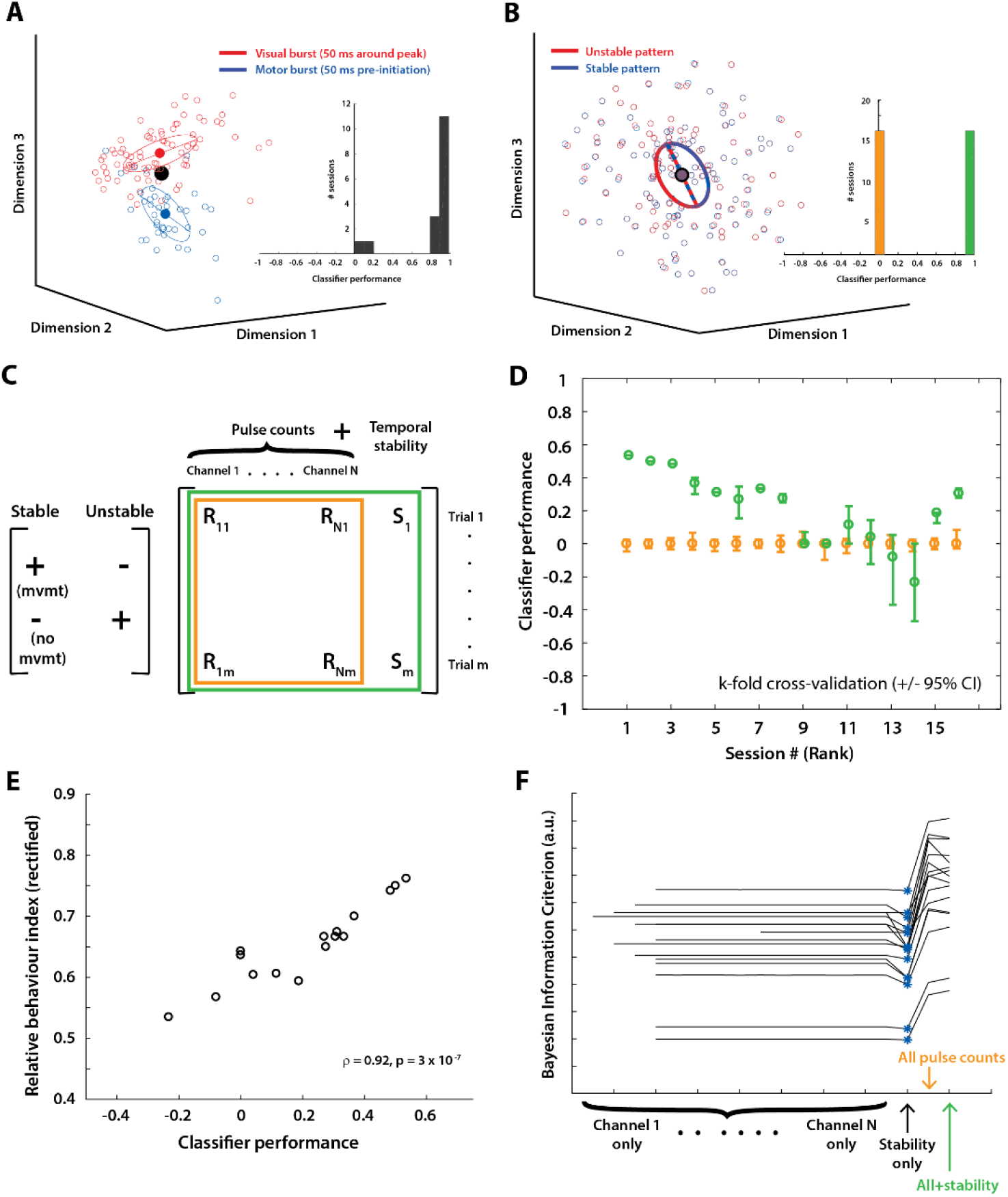
Linear discriminability of population activity and microstimulation patterns. **A**.Three-dimensional factor analysis (FA) projection of population activity states during the visual and premotor bursts (red and blue points, respectively; 50 ms around peak of visual burst, 50 ms before saccade initiation in premotor burst. The dimensions were chosen arbitrarily to illustrate the separation of the two states. The solid lines and ellipses show the axes of maximum variance and covariance ellipses for each cluster, respectively. The histogram in the inset shows the performance of a linear discriminant analysis (LDA) classifier on labelling visual and premotor states on individual sessions (perfect classification is 1, chance is 0). **B**. FA projection of population pulse states for stable and unstable microstimulation patterns (blue and red points, respectively). The projection dimensions were chosen arbitrarily, but no view showed meaningful separation between the two clusters. The histogram in the inset shows LDA performance on classifying stable and unstable stimulation patterns, based on population pulse states alone (orange bars), or with a combination of population states and temporal stability (green bars). Note that perfect classification is expected by definition in the latter case. **C**. Schematic of the trials and data matrix used for classification of stimulation-evoked movement occurrence. Left - Trial pairs in which only one of the stable or unstable pattern evoked a movement. Right - The classifier was first trained on pulse counts alone (part of the matrix highlighted by the orange boundary – match color to the next panel). Separately, the classifier was also trained on an additional dimension of temporal stability values (full matrix highlighted by the green boundary). **D**. LDA classification performance on discriminating trials in which stimulation evoked a movement from those in which it did not. The orange points are with the classifier trained solely on the population pulse states. The green points are with the addition of temporal stability. For most sessions, addition of temporal information increased classification performance (p < 0.01, one-tailed t-test of a difference between the crossvalidated performance distributions with and without addition of temporal stability). Sessions are sorted by the order determined in Figure 3C based on relative likelihood of evoking a movement. **E**. Scatter plot showing correlation between classifier performance and behavior (values of blue points in Figure 3C rectified around 0.5). The classifier’s discriminative ability predicted the relative behavioral effect size between stable and unstable patterns. **F**. Model selection for discriminating between trials in which a movement occurred versus those in which it did not occur. Bayesian information criterion (BIC) traces as a function of model type for individual stimulation sessions show that the lowest BIC (blue asterisks) was attained for the stability-only model for every session.

Next, we used data from the patterned microstimulation experiments to disambiguate between the optimal subspace and temporal stability frameworks. Using a representative low-dimensional FA projection to differentially visualize stable and unstable stimulation patterns was infeasible because all the electrode sites contributed equally to the patterns, due to the random assignment of pulse rates across sites on different trials. From the example 3-dimensional projection in Figure 4B, it is evident that the stable and unstable patterns are indiscriminable in state space. Indeed, an LDA classifier failed to discriminate between stable and unstable patterns based on population pulse states alone, never yielding above-chance classification (Figure 4B, orange bar in histogram inset). However, an LDA model that also included stability information was able to classify perfectly, reaffirming the difference in temporal structure (by design) between the two sets of patterns (Figure 4B, green bar in inset; also see right matrix in Figure 4C). These results indirectly provide a key piece of evidence in support of the notion that the brain in fact uses temporal information, since the evoked behavior reflected the difference between stable and unstable patterns while a static linear readout of population states did not.

The next set of analyses examined the effect of stimulation pattern on behavior. We focused on the trial pairs in which only one of stable or unstable pattern evoked a movement (left matrix in Figure 4C), since these are likely to provide insight into the discriminatory role played by temporal structure. We trained another LDA classifier to discriminate between trials in which stimulation evoked a movement and those where no movement was evoked. When using the pulse states alone, the linear decoder was unable to determine whether a movement was evoked on a given trial with above-chance accuracy (orange points in Figure 4D). Since the stable-unstable pairs were matched in terms of whether a movement was evoked or not for this subset, this readout of the population pattern was independent of its temporal structure. Crucially, and in contrast, when the population pulse states were supplemented with stability information (added as a native dimension in the input to the decoder, see green matrix in Figure 4C), classifier performance increased to reflect the broad trend in relative movement likelihood (compare blue points in Figure 3C with green points in Figure 4D). The correlation between performance of the full LDA classifier and the behavioral effects of stimulation is better highlighted in Figure 4E, i.e., the brain’s ability to extract temporal structure to generate movements is reflected by a model that utilizes this temporal information. We also independently evaluated whether temporal stability was more informative than each and all pulse count dimensions by performing model selection analysis with a logistic regression fit to the movement occurrence data used for the previous analysis. For every stimulation session, the best model (one with the lowest Bayesian Information Criterion or BIC) was one that only had the temporal stability dimension (blue asterisks indicating minima in BIC curves in Figure 4F). Together with the effect of dispersion seen earlier (Supplementary Fig. 6), these analyses reveal two independent contributors to movement initiation in SC – a static “laminar” code, presumably related to the optimal or potent subspace, that may be a function of the distribution of preferred stimulation sites, and a dynamic “temporal” code, where stability of the population pattern controls movement initiation *even* under matched state space conditions.

We also tested how the temporal stability hypothesis was related to modified threshold-based mechanisms that rely on pooling the activity of a correlated ensemble of neurons (*9, 32*). In this framework, correlations in the accumulation rates of neurons influence movement initiation (reaction times) under a given pooling scheme, with higher correlations leading to shorter reaction times. Spike count correlations between SC neurons in the visual and premotor epochs (for details, see Methods) were inconsistent with this notion – the correlations during the visual epoch were slightly, but significantly, higher compared to the premotor epoch (Supplementary Fig. 7A, mean difference between visual and premotor epoch correlations = 0.0167 > 0, p = 2.5E-6, one-tailed t-test). We also computed pulse count correlations for the stable and unstable stimulation patterns and found no difference (Supplementary Fig. 7B, median difference in correlations between conditions = 4.11E-4, not significantly different from 0, p = 0.279, Wilcoxon signed-rank test), despite the significant difference in their effects on behavior. Thus, the temporal stability framework is, to a large extent, independent of extant models of movement initiation.

### Biophysical models with synaptic facilitation can decode population temporal structure

Finally, we sought to identify a mechanism by which downstream neurons could discriminate between stable and unstable population codes. We modelled the decoder as a spiking neuron that receives population inputs through its network of dendrites (Figure 5A). The decoder can be thought to represent neurons in the pons that receive and integrate SC inputs and burst for saccades (*27*). To mimic the potent inhibitory gating (and disinhibition during saccades) provided by the omnipause neurons (OPNs) on the burst neurons, we also included a spiking disinhibitor unit with reciprocal inhibitory connections with the decoder. The disinhibitor also received both excitatory (*33*) and inhibitory (*34*) inputs from the same population as the decoder, creating a balance between excitation and inhibition. This balance must be overcome by some property of the input activity in order to produce a saccade. How might the population pattern be converted into a signal that initiates a movement only when the inputs are stable? Conceptually, the decoder should have a mechanism to keep track of the short-term history of population activity, use this history to evaluate temporal structure, and respond selectively when the activity pattern is deemed stable over the time scale of integration. We incorporated these heuristic requirements by using state-dependent modulation of input-evoked excitatory post-synaptic potentials in the decoder, i.e., inputs arriving when the local post-synaptic potential was depolarized caused a higher subsequent change in the potential (*35, 36*) (Figure 5B).

**Figure 5.**
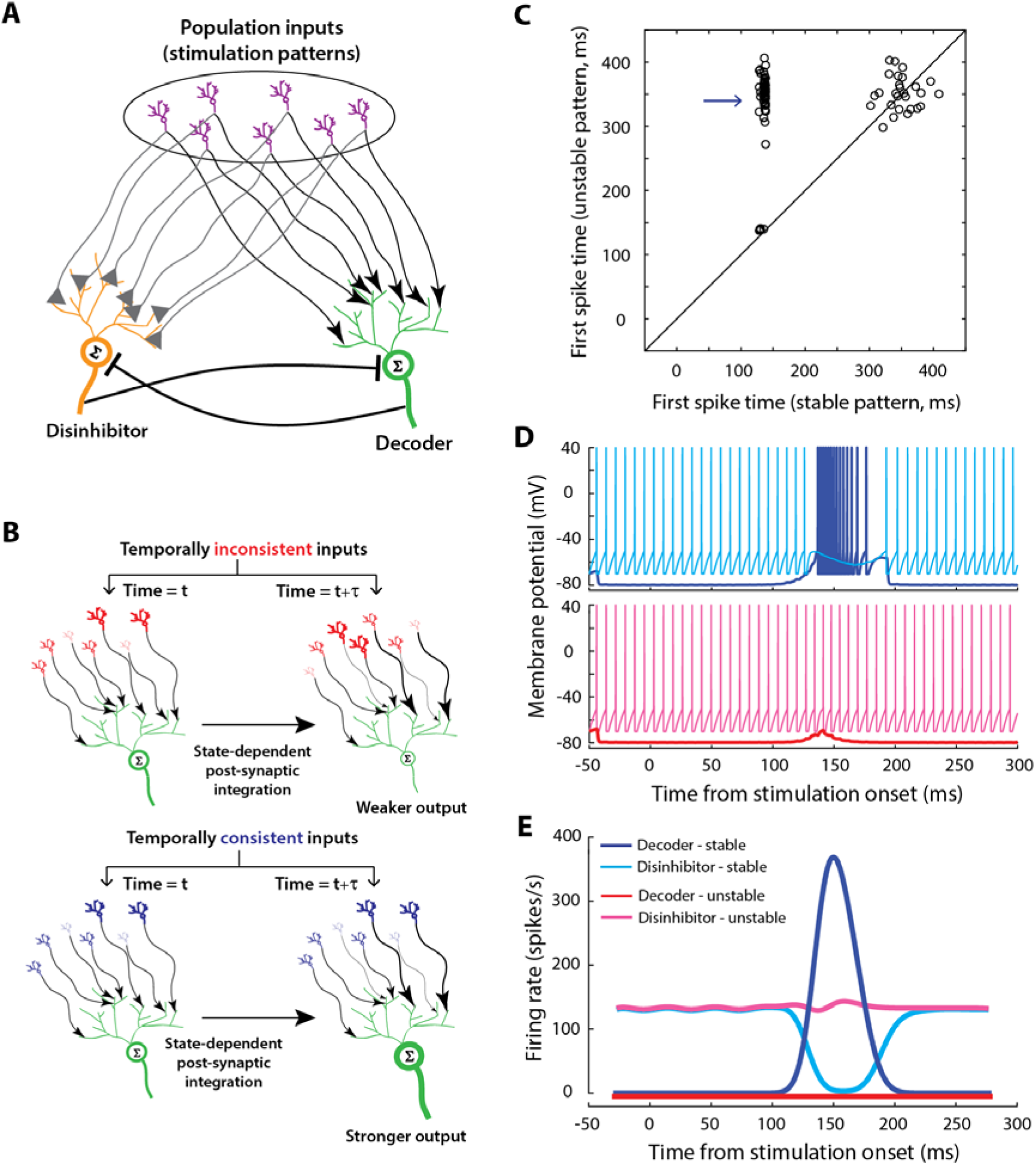
Spiking neuron model with dendritic processes can discriminate population temporal structure. **A**. Schematic of the model architecture. The decoder (green neuron), representing high frequency burst neurons in the brainstem, receives population inputs (output of purple neurons, or, in this case, microstimulation pulses) through its dendritic network (excitatory inputs, represented by the arrowhead terminals). The disinhibitor (orange neuron), representing pontine omnipause neurons, also receives inputs from the same population (both excitatory and inhibitory(*33, 34*), represented by the triangular terminals), in addition to a constant current that produces tonic firing activity (not shown). The decoder and disinhibitor are both modelled as leaky integrate-and-fire spiking neurons and mutually inhibit each other (flat terminals). **B**. Schematic of the model heuristic. Each input neuron’s activation level is represented by its size and thickness. The effect of an input spike at the local dendritic site (i.e., excitatory post-synaptic conductance changes) is represented by the thickness of the arrow. Two time points are shown for illustration (left column – arbitrary initial point, right column – subsequent time point). In the top row, unstable inputs lead to a scenario where the post-synaptic conductances are no longer aligned to the strong inputs that initially created them, whereas in the bottom, stable inputs lead to matched strong input and post-synaptic conductances, resulting in stronger accumulation and firing. **C.** Scatter plot of first spike latencies in the decoder (putatively representing saccade initiation) for stable versus unstable inputs from matched pairs. The input duration was 150 ms and thus only latency values <150 ms correspond to actual first spikes produced by the decoder. To facilitate comparison with Figure 3B, latency values were assigned randomly for trials in which the decoder did not generate any spikes, sampled from a distribution with an arbitrarily selected mean of 350 ms. Note the occurrence of a number of trial pairs for which only the stable input causes the decoder emit spikes (blue arrow). **D.** Simulated membrane potential of the decoder (blue and red traces) and disinhibitor (cyan and magenta traces) for an example matched trial pair with stable (top row) and unstable (bottom row) input patterns. **E.** Average activity of the disinhibitor (cyan and magenta traces, stable and unstable inputs, respectively) and decoder (blue and red traces, stable and unstable inputs, respectively) for all trial pairs in which only the stable input produced a spike/movement (i.e., trial pairs indicated by the arrow in panel D).

We simulated the model with the stable and unstable patterns from the microstimulation experiments as inputs, since they readily offered mean-matched input sequences differing only in temporal structure. We characterized the efficacy of integration by the decoder as the time of first spike in the burst, a proxy for movement initiation latency. Although there were several pairs of trials for which both or neither the stable and unstable inputs caused the decoder to spike (Figure 5C), there were a number of pairs for which only the stable input pattern was decoded (arrow in Figure 5C). To facilitate visualization and comparison with experimental data (Figure 3B), latency values greater than the stimulation duration were randomly assigned when the decoder wasn’t recruited; this provides a visualization of the number of trials where no movement was evoked by either pulse train of the pair. Figure 5D shows the membrane potential output of the disinhibitor (cyan and magenta traces) and decoder (blue and red traces) units for a pair of stable (top row) and unstable (bottom row) input patterns from this subset of trial pairs. The disinhibitor maintained a tonic firing rate throughout the trial when the inputs were unstable, and sharply reduced its activity at some point during stable stimulation (Figure 5E - magenta and cyan traces, respectively). This latter disinhibition on stable trials was in anti-phase relationship with the bursting exhibited by the decoder, which was absent on unstable trials (Figure 5E - blue and red traces, respectively). See Supplementary Fig. 8 for membrane potential waveforms when both or neither the stable and unstable patterns recruit the decoder unit. The firing rate profiles of the decoder and disinhibitor units share a striking resemblance to medium-lead burst neurons and OPNs, respectively, in the pons (*37*).

Thus, a relatively simple, yet biophysically realistic, module seems capable of discriminating between population inputs based solely on their temporal characteristics, offering a putative mechanism by which downstream networks could use temporal information to make decisions about movement generation. The model simulations predict that temporal structure will impact integration near activation threshold, but a surplus of spikes in an unstable input may still be integrated well enough to produce a movement, consistent with the observations in the patterned microstimulation experiment. Another testable prediction is that disinhibition of the gaze control circuitry by silencing the OPNs should diminish the effect of temporal structure in determining movement initiation, producing movements for both stable and unstable inputs. We also found that a firing rate-based accumulator model with short-term synaptic plasticity produces comparable results (Supplementary Fig. 9), demonstrating the flexibility of various biophysical mechanisms in their ability to decode temporal structure and hence, the model-independence of this result.

## DISCUSSION

Neurons in premotor structures are constantly bombarded with information from thousands of presynaptic neurons that are active during sensorimotor processing. It is unclear how activity relevant for movement initiation is discriminated from activity related to other processing. Extant models of movement generation that rely on firing rate, including rise-to-threshold (*6*), inhibitory gating (*37*), and dynamical switches at the population level (*12,13*), leave certain explanatory gaps unfilled. A canonical model of movement initiation, especially in the oculomotor system, is threshold-based gating(*6*) (Figure 6A-B). Current knowledge points to a role by the OPNs in defining the threshold and controlling saccade initiation (*6, 38*). However, thresholds vary across behavioral paradigms (*8, 39*), raising the question of how the threshold is set in a particular condition. Furthermore, evidence that the threshold changes during the course of a trial purely based on OPN activity is limited; OPN activity increases transiently during the visual burst but only by an additional spike or two and only in a small subset of neurons (*38*). Critically, the existence of trials in which the population activity during the visual response exceeds premotor activity strongly reduces the likelihood of thresholding operating as a singular mechanism.

**Figure 6.**
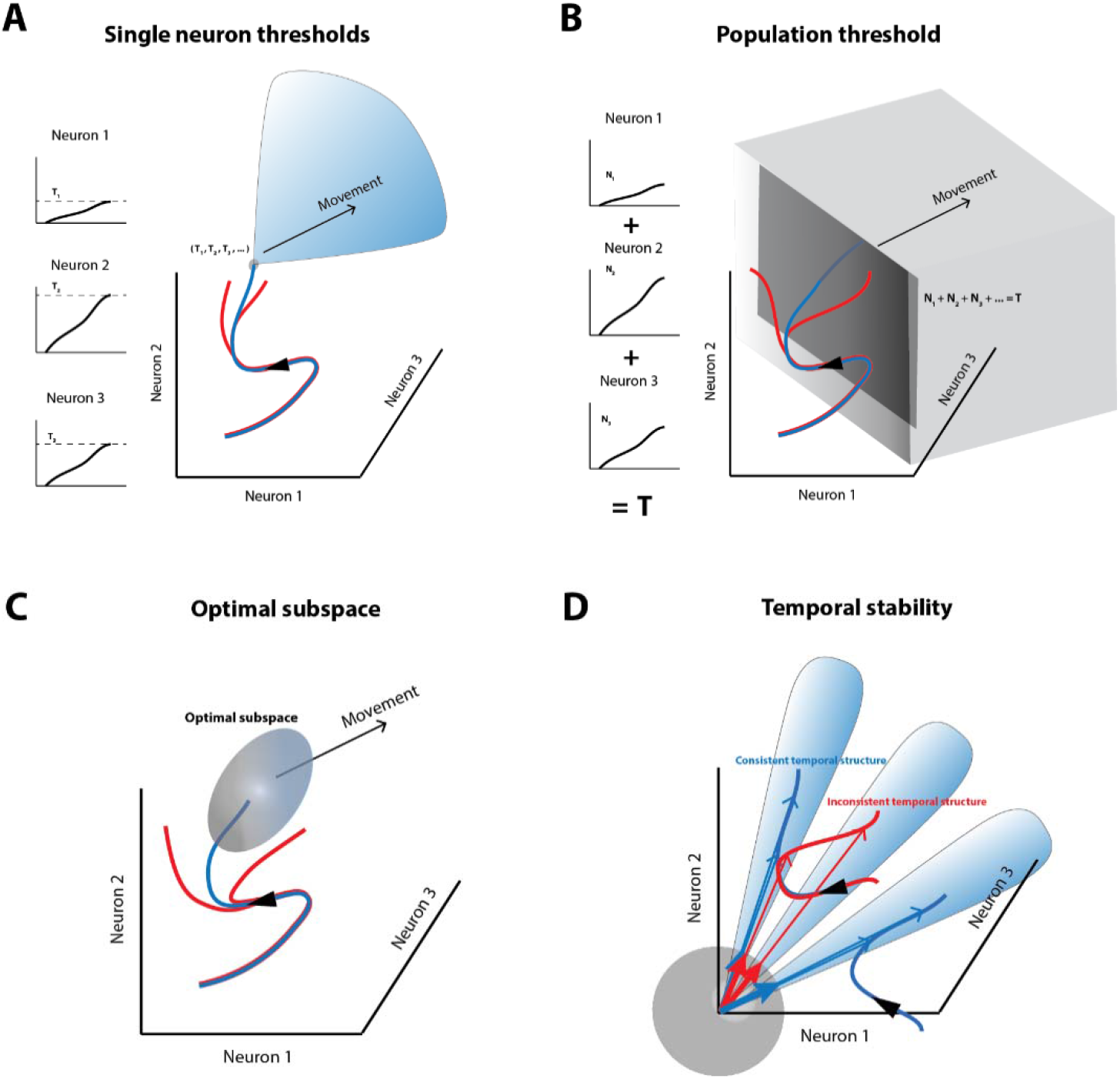
Summary of models of movement preparation in a state space framework. **A** and **B**. Multi-dimensional representations of the threshold hypotheses. Panel A depicts single neuron thresholds, i.e., the activity of each neuron must rise to a fixed threshold at the same time in order to initiate a movement. Fixed thresholds for each neuron are equivalent to a point (or a small region, represented by the gray sphere) in neural state space. Activity profiles that meet this criterion (e.g., blue trajectory) and beyond (blue region) lead to movement generation while those that don’t meet this criterion (red traces) do not result in a movement. Alternatively, activity may need to cross a fixed threshold at the population level (sum of neurons’ activities = constant, i.e., an n-1 dimensional hyperplane, dark gray plane), depicted in panel B. **C**. The optimal subspace hypothesis offers more leeway by allowing neuronal activity to reach a relatively larger region of population state space (blue trajectory and gray ellipsoid) over a period of time in order to signal the initiation of a movement. Activity trajectories that evolve outside this “optimal subspace” (red traces) do not lead to a movement. **D**. The temporal stability hypothesis suggests that a burst of neural activity that is consistent over time in state space, i.e., an activity trajectory that points in the same direction (blue trajectories and vectors) is likely to be interpreted as a movement command by a decoder, while high-frequency activity that is inconsistent (fluctuating directions, red trajectories and arrows) is not. It does not matter where in state space the activity happens to evolve – a different subpopulation of neurons could be active, but as long as they are pointing in the same direction (i.e., evolving within one of the blue sectors), it will lead to a movement. The gray sphere around the origin represents a unit hypersphere for visualization of back-projected unit vectors.

An extension of simple thresholding is a mechanism based on the pooled activity of neurons with varying degrees of correlation in the population (*9*). However, we show that population correlation statistics are not necessarily related to temporal structure or saccade initiation (Supplementary Fig. 7). A related alternative explanation is that a movement is initiated specifically when movement-only neurons are active, but most neurons in sensorimotor structures like SC and FEF span a continuum between visuomovement activity and movement activity (*1, 31, 40*). In fact, antidromically identified tectoreticular neurons that presumably drive saccade generation exhibit both visual and premotor bursts(*4*). Moreover, as seen earlier, SC neurons with the highest correlation to movement velocity tend to also have strong visual responses (Supplementary Fig. 1). Finally, the FEF also projects to the premotor circuitry in the brainstem (*25*). Our pseudo-population analysis revealed similar temporal stability waveforms for both regions considered individually (Figure 2A, colored traces) as well as collectively (Figure 2A, black trace). Incorporating the result that impairment of either region alters saccade kinematics produced by the other (*41, 42*) together with these results suggests that both regions operate cooperatively to initiate saccades.

Other mechanisms, primarily based on static weighted linear readouts of population activity, have been proposed for how movement generation unfolds in the skeletomotor system. These models posit that muscles are recruited and a movement is initiated when neural activity traverses an optimal or motor-potent region of the population state space (*12, 13*) and is inhibited otherwise, e.g., during movement preparation (*13, 14*) (Figure 6C). While the potent- and null- subspaces hypothesis certainly seem to be consistent with recorded neural activity in SC (Figure 4A), they cannot readily account for the observation that neck (*43*) and upper limb (*44*) muscles are recruited time-locked to the visual target, as proxied through electromyography. We instead show that temporal stability plays an independent role in determining movement initiation, even when the putative potent subspace is matched in a causal manipulation (Figure 4D, green symbols). We reason that for a decoder downstream, it is important to ensure the stability - or consistency over time - of the input code while processing it in order to influence the motor output. Population activity that creates a high firing rate drive but is unstable is also likely unreliable and must have a reduced efficacy of triggering a movement. This requirement is critical especially in the case of ballistic movements such as saccades, where the ability to reverse the decision once the movement has been initiated is limited.

Thus, we present a novel mechanism, wherein both high firing rate and consistent temporal structure are necessary conditions for movement initiation, allowing for population trajectories that traverse away (increasing overall rate) but within hypersectors that fan out (stable temporal pattern) from the state space origin (Figure 6D). Note that this model does not preclude population activity from traversing a localized “optimal” region of state space, possibly due to hardwired network constraints. Crucially moreover, the proposed mechanism is also consistent with classic population vector decoding schemes (*45, 46*), since any weighted population vector readout during the sensory and movement epochs will produce decoded movement vectors that are similar given a sufficiently large integration window. This allows for the same physical movement to be planned and executed by a given population of neurons in the labelled line sense, a mechanism that is precluded by weighted neural readout mechanisms of movement preparation and generation, such as the potent/null space models.

Our findings are closely related to the premotor theory of attention (*47*) and offer a way to reconcile the attention-intention debate. They could also account for the mirror-like activity recorded during both action observation and execution in neurons known to project directly to motoneurons in the spinal cord (*48*). In both cases, the same neuronal population represents two distinct signals that serve different functional roles. The results presented here suggest that this multiplexing ability may be provided by the distinct temporal structures of population activity patterns. Indeed, intracellular recordings have demonstrated that visual stimulation drives cortical networks into an asynchronous state (*49*), which may be a critical requirement for sensory processing.

Additionally, the differential effects of stable and unstable population stimulation patterns are reminiscent of the desynchronizing effects of patterned deep brain stimulation recently developed for the treatment of Parkinson’s disease (*50*). In this so-called “coordinated reset” approach, sequential, non-overlapping spatiotemporal sequences are more effective at resetting the affected neuronal population from the pathological synchronized state to an asynchronous state, enabling better movement control (*51*).

Previous studies have looked at the role of precise coordination in the timing of incoming spikes in the transmission of information and the efficacy of driving the recipient neuron (*20, 21*). However, input firing rates may vary greatly across the population, limiting the ability to compare to spike times. Our study proposes a mean-field equivalent to the spike-based temporal or correlation code (*22, 52*) by looking at the temporal structure of population firing rates, thus tying together the notion of rate, temporal, and population codes. We suggest that temporal structure of population activity is critical to understanding movement generation as well as, more broadly, neuronal communication and its relationship to behavior.

## METHODS

### Subjects and Surgical Procedures

All experimental and surgical procedures were approved by the Institutional Animal Care and Use Committee at the University of Pittsburgh and were in compliance with the US Public Health Service policy on the humane care and use of laboratory animals. We used three adult rhesus monkeys (*Macaco mulatto*, 2 male and 1 female, ages 6 -12) for our experiments. Both SC and FEF were recorded in monkeys BB and BL whereas only SC was recorded in monkey WM. Under isoflourane anaesthesia, recording chambers were secured to the skull over craniotomies that allowed access to the SC and FEF. In addition, posts for head restraint and scleral search coils to track gaze were implanted. Analgesic and antibiotic treatment protocols were followed after surgery. Post-recovery, the animal was trained to perform standard eye movement tasks for a liquid reward.

### Visual stimuli and behavior

Visual stimuli were displayed either by back-projection onto a hemispherical dome or on a LED-backlit flat screen monitor. Stimuli were white squares on a dark grey background, 4×4 pixels in size and subtended approximately 0.5° of visual angle. Eye position was recorded using the scleral search coil technique or using an EyeLink 1000 eye tracker, both sampled at 1 kHz. Stimulus presentation and the animal’s behavior were under real-time control with a Labview-based controller interface(*55*). All monkeys were trained to perform standard oculomotor tasks. In the delayed saccade task (Figure 1A), the monkey was required to initiate the trial by aligning gaze on a central fixation target. Next, a target appeared in the periphery, but the fixation point remained illuminated for a variable 500-1200 ms, and the animal was required to delay saccade onset until the fixation point was extinguished (GO cue). The gap task (Figure 2D and Supplementary Fig. 5A) was used for express saccade analysis and for the patterned microstimulation experiments. In this task, initial fixation on a central target was followed by a gap period (200-300 ms) during which the fixation point disappeared, while the animal was required to maintain fixation at the now vacant location. On stimulation trials, microstimulation pulses were delivered 100 ms into the gap period and window constraints were relaxed to allow for the stimulation-evoked movements. This was followed by the appearance of a saccade target in the periphery which was also the GO cue for the animal to make a target-directed saccade. All animals performed these tasks with >95% accuracy. Incorrectly performed trials were removed from further analyses. The tasks were occasionally interleaved with visual search paradigms used for a different study.

### Experimental sessions

We used data from several different types of sessions in this study. Each session was either a laminar recording session (n = 20 sessions), a single-unit recording session (n = 108 sessions), or a patterned microstimulation session (n = 16 sessions). The laminar and single-unit recording sessions employed mostly the delayed saccade task with a few microstimulation trials to verify electrode presence in SC (or FEF) and estimate the location on the topographic map. The patterned microstimulation sessions were restricted to the gap task, with stimulation trials and non-stimulation trials interleaved.

### Laminar recordings and single-electrode electrophysiology

During each laminar recording session, a linear microelectrode array (LMA, AlphaOmega, Inc.) or a Plexon V-probe (Plexon, Inc.) was inserted into the SC chamber using a hydraulic microdrive. The arrays had either 16 or 24 contacts, with inter-contact spacing of 150 μm or 175 μm, respectively. Neural activity was amplified, digitized and recorded using the Grapevine Neural Interface Processor (Ripple, Inc.) and visualized using the associated Trellis interface. Neural activity was band-pass filtered between 500 Hz and 5 kHz to record spiking activity and between 0.1-250 Hz to record local field potentials for another study. Approach towards the SC surface was identified by luminance-based visual modulation of activity in the lowermost channels, after which the electrode was driven down a further 2-3 mm until known hallmarks of SC activity were observed in a majority of the channels. The presence of several contacts in the intermediate layers was further confirmed by the ability to evoke movements with single-channel microstimulation at these contacts. For the initial subset of experiments, a tungsten microelectrode was lowered into the FEF or SC. Neural activity was amplified and band-pass filtered between 200 Hz and 5 kHz and fed to a digital oscilloscope for visualization and spike discrimination. A window discriminator was used to threshold and trigger spikes online, and the corresponding spike times were recorded. Both SC and FEF were confirmed by the presence of visual and movement-related activity as well as the ability to evoke fixed vector saccadic eye movements at low stimulation currents (20-40 μA, 400 Hz, 100 ms). Before beginning data collection for a single or laminar recording site, the neural response field was roughly estimated. Since the electrode track was directed orthogonal to the SC surface, most neurons preferred similar visual response fields and/or movement vectors. In most data collection sessions with either electrode, the saccade target was placed either in the neurons’ response field or at the diametrically opposite location in a randomly interleaved manner. In addition, stimulation-evoked saccades were recorded to identify the response field centers (or “hotspots”) for the cells recorded during that session. For recordings in rostral SC, stimuli were presented at one of four locations at an eccentricity sufficient to induce a reduction in activity during the large amplitude saccade.

### Data analysis and pre-processing

Data were analyzed using a combination of in-house software and Matlab. Eye position signals were smoothed with a phase-neutral filter and differentiated to obtain velocity traces. Saccades were detected using standard velocity criteria (30-50°/s). The animal was considered to be maintaining fixation if the gaze remained within a 2-3° window around the fixation target. We also detected any microsaccades that occurred during the delay period in each trial by using a velocity criterion based on the noise in the velocity signal for that trial. Only one of the two monkeys (WM) in whom we recorded neural activity in the rostral SC made sufficient number of microsaccades to permit further analysis.

Raw voltage signals filtered to spiking frequencies (500 Hz – 5 kHz, see above) were passed through a threshold criterion to obtain spike times corresponding to multi-unit activity (MUA). Spike density waveforms were computed for each neuron (or multi-unit activity cluster) and each trial by convolving the spike trains with the causal portion of a Gaussian kernel (width = 5 ms; in some instances, we used 10 ms for display purposes). Data from the patterned microstimulation experiments (n = 16 sessions) were also subjected to this procedure to compute pulse density waveforms. For the laminar recordings, we analyzed channel activity on single trials independently. Because direct kernel-based estimation of firing rates from binary spike trains on single trials can be noisy, we first computed the inverse of the inter-spike intervals as a measure of the firing rate prior to convolution with the smoothing kernel. All units that crossed the spiking threshold of 6σ were considered for further analyses, regardless of their characteristics (i.e., number of neurons/multi-unit clusters matched number of channels for these datasets). Only sessions which had at least 6 channels with typical visuomovement activity patterns were considered for further analysis, resulting in 16 laminar recording sessions (monkey BB – 104 channels/6 sessions, monkey BL – 176 channels/10 sessions). For single-electrode recordings, the spike densities from condition-matched trials were randomly combined to form a pseudo-population dataset (see next section). Neurons were classified as visuomovement neurons if the spike density was significantly elevated above baseline during the visual epoch (50-200 ms following target onset) and during the premotor epoch (50 ms before and after saccade onset). This resulted in 60 caudal SC neurons and 30 FEF neurons. In addition, rostral SC neurons were defined as fixation-related if the activity during the premotor epoch of large saccades was significantly reduced below baseline (36 neurons). A subset of these neurons also elevated their discharge around the onset of microsaccades (13 neurons). To minimize the effect of noise in the spike density waveforms due to insufficient number of trials in our analysis, we used only neurons which had at least 10 trials for a given condition. This was not a factor in most cases (we typically had 50-100 trials). For the gap task analysis, we used neurons from sessions that had at least 10 express saccade trials, defined as the set of trials in the first peak of the saccade reaction time (SRT) histogram (SRT < 150ms). Regular saccade trials comprised the second peak (SRT > 150ms).

To verify that spike isolation quality did not have an undue effect on our analyses, we also repeated the main analysis from Figure 1 on spike sorted data. Spike sorting was performed using Kilosort2, an algorithm developed for dense laminar probe recordings such as those with Neuropixels probes (https://www.ucl.ac.uk/neuropixels/spike-sorting). The results of sorting were further manually inspected to remove obvious misclassification errors. Sorted spike unit times were converted to spike density waveforms using the same procedure as for MUA laid out above. When multiple units were detected in a single laminar electrode channel, they were treated as distinct components of the population vector corresponding to that session. The sorting produced 391 units over 280 channels across 16 sessions (session counts – 12, 22, 23, 18, 24, 13, 40, 20, 27, 40, 23, 15, 29, 32, 29, 24 units). The results of the analysis on spike sorted data strongly resembled the main temporal stability results from MUA (compare Supplementary Fig. 3 and Fig. 1). Therefore, we restricted all analyses in the rest of the paper to MUA, especially since spike sorting has been recently also found to have a negligible effect on population-level analyses(*23*), and unsorted MUA may be more representative of the overall input processed by downstream brain networks.

### Estimating population dynamics from laminar and single-unit recordings

We define **R**(t) as a population activity vector:

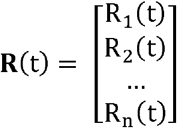

where **R**(t) represents the instantaneous activity at time t as a point in an *n*-dimensional space, *n* is the number of neurons, and R_i_(t) is the spike density function of the *i*^*th*^ neuron. The curve connecting successive points over time is the neural trajectory that describes the evolution of population activity. For laminar recording data, a neural trajectory was determined directly for each trial. Analysis of single electrode data relied on pseudo-population analysis, for which a surrogate dataset was created with individual trials drawn from each single neuron session. To ensure that these “pseudo-trials” were matched in relevant metrics, we first binned trials from each session into quartiles based on saccade reaction time and length of the delay period (the latter only for the delayed saccade task). A pseudo-trial was then created by drawing randomly from matched quartiles for each session, and this process was repeated 1000 times to create a full surrogate dataset from which pseudo-population trajectories were computed for individual pseudo-trials. Since SC and FEF neurons have fairly broad visual and movement fields, many neurons in each region contribute to the active population for any given visual stimulus or saccade(*56*), respectively. Our neurons were sampled roughly (but not exactly) around the hotspot of the active population for a given session. Therefore, the pooled data from individual sessions can be thought of as an approximation of the population mound active for an arbitrary location/RF in the visual field on any given trial. Many recent studies have reconstructed such pseudo-populations from sequentially recorded neurons and found comparable properties from simultaneous and serial recordings and found comparable properties from simultaneous and serial recordings(*13, 57, 58*). Indeed, this is also expected of our pseudo-population under the assumption of isotropy – that each neuron’s contribution to its respective local active population is similar regardless of the locus of the population, and consistency between the results we observed in the laminar dataset and pseudopopulation confirms this. To better demonstrate this, we estimated the location of a given neuron in the active pseudo-population as follows (we use SC for illustrative purposes because of its convenient topography). We mapped the target location onto the SC map and used this locus as a representation of the center of the active population for that session, and used the stimulation-evoked saccade vector to identify the location of that neuron on the SC. We referenced the mapped locus of each target location to a single location to create an active pseudo-population and translated the neuron locations relative to this population center (Supplementary Fig. 4A). All mathematical equations for transforming between visual and SC tissue coordinates have been defined previously(*59*). Inclusion of stimuli/saccades in the anti-preferred RF of the neurons allowed us to also estimate a pseudo-population of neurons in the ipsilateral SC. To complete the representation of activity across the SC topography, we also recorded from and included in our analyses neurons in the rostral portion of SC, which are active during fixation(*26*), burst during microsaccades(*28*), and are suppressed during large saccades(*26*).

### Temporal stability analysis

To assess temporal stability, we first normalized the population trajectory **R**(t) by its Euclidean norm (||**R**(t)||), equivalently its magnitude, at each time point to yield 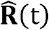:

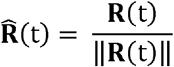

The normalized trajectory can be visualized as a unity length population vector that points in an *n*-dimensional direction at each instant in time. That is, while **R**(t) is free to traverse the n-dimensional activity space, 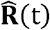 is constrained to the surface of an *n*-dimensional hypersphere (Figure 1C). Temporal stability or consistency of the evolving population was then quantified by the dot product of two timeshifted unity length vectors:

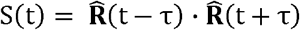

The stability metric (S(t)) tracks the running similarity of the normalized trajectory separated in time by 2 τ ms. Crucially, the normalization constrains S(t) between 0 and 1. Thus, if S(t) → 1, the population activity is considered stable since the relative contribution of each neuron is consistent. If S(t) « 1, the population activity is deemed unstable because the relative contribution is variable. If the neural trajectory is not normalized, the dot product quantifies similarity across the vectors’ magnitude and direction. It roughly mimics the quadratic of the firing rate (exactly so for *τ* = 0). When the vector direction remains constant, the dot product yields no additional information than that already present in the firing rate. In contrast, the normalization scales the neural trajectory and thus always has unity magnitude. It neither alters the relative contributions of the neurons nor compromises the vector direction. The dot product therefore performs an unbiased evaluation of stability based only on vector direction and is an estimate of the fidelity of the population code modulo a multiplicative gain factor. Intuitively, the stability measure is analogous to the correlation between the neurons’ activities at two different time points. We chose the dot product, however, because of its interpretability as a measure of pattern similarity in *n*-dimensional activity space.

We assessed the significance of the stability profiles in two ways. First, we compared the across-trial distribution of stability values in appropriate windows around the visual burst and during the premotor epoch for each session (Figure 1F). For this analysis, the stability values were averaged in a 20 ms window preceding the peak of the visual burst for the visual epoch and in a 20 ms window preceding the saccade initiation time (15 ms before saccade onset) in the premotor burst for the premotor epoch. Second, for each trial, we randomly shuffled the activities of various neurons (channels for laminar recordings) at each time point (dark gray trace, Supplementary Fig. 2A, bottom row), or randomly shuffled the time points at which the population patterns occurred during the trial (light gray trace, Supplementary Fig. 2A, bottom row). The former shuffle retains the average firing rate at each instant and the latter shuffle retains the overall population correlation structure but both shuffles remove any temporal correlation in the firing rate across neurons. We performed multiple such shuffles and recomputed temporal stability for each instance, followed by across-trial averaging, baseline correction, and across-session averaging to obtain the distribution of stability profiles expected to occur by chance if the neurons’ activities were uncoordinated.

To mitigate the effect of potentially variable visual response onset latencies (on different trials) on the trial- and session-averaged mean temporal stability profiles in the visual epoch, we also performed the averaging after realigning the data to the peak visual response on individual trials (Figure 1E). In order to do this, the time of peak visual burst was identified from the population mean (average across channels) firing rate on individual trials and was used to align the traces across trials before computing the trial average.

For the mean-matched control in Figure 1D&E, we performed the temporal stability analysis only on the subset of trials in which the peak of the population visual response was greater than or equal to the population activity at the time of saccade initiation, after accounting for an efferent delay. We chose 15 ms before saccade onset as the latest time at which SC activity could influence saccade initiation, because it is consistent with a range of putative efferent delays reported using various approaches(*6, 16, 17*).

### Patterned microstimulation

The patterned microstimulation experiments allowed us to causally evaluate and compare the temporal stability model to other models of movement initiation. Given the ability to control the delivery of individual stimulation pulses to each contact on the laminar electrode, we used stimulation patterns with specific spatiotemporal features to evaluate their relative efficacy in evoking movements. We designed these patterns as follows. All individual stimulation pulses were biphasic with a leading cathodic phase, monophasic pulse widths of 200-267 μs, and an interphase interval of 100 μs. Our first step was to determine the appropriate range of current intensities and frequencies for each experiment. During each session (i.e., for each electrode penetration into SC, total n = 16 sessions), we first stimulated from individual contacts with standard suprathreshold parameters (40 μA, 400 Hz) to identify the set of contacts that elicited a movement at a sub-50 ms latency. We restricted the experiment to these contacts for the rest of that session. We then stimulated simultaneously across all these contacts at the same current and frequency, starting from 4 μA, 100 Hz, stepping up by 1 μA and 50 Hz, to determine the threshold for evoking a movement with multi-channel stimulation. The threshold, defined as the current/frequency combination for which stimulation generated movements on approximately half the trials, ranged from 5-12 μA and 150-200 Hz across sessions. For each session, we used the metrics of the saccades evoked at these threshold parameters from constant frequency stimulation in lieu of “fixed vector saccades” as a control against which to compare the patterned microstimulation results (Supplementary Fig. 5D&E).

Once we identified the threshold parameters, we designed stimulation patterns at that current intensity but with time-varying frequencies intended to simulate a burst of activity. Each stable pattern was created with a linearly decreasing inter-pulse interval (IPI) sequence for one contact (to simulate a burst) and scaling the IPIs for other contacts by a uniformly spaced factor (see below). The 150 ms duration burst was composed of a flat baseline (40 ms), a rising phase (80 ms), and a falling phase (30 ms). The peak frequency for individual channels ranged uniformly between 20 Hz and 400 Hz with the mean peak frequency across contacts pegged to the threshold frequency determined from the constant frequency multi-channel stimulation in an earlier step. At these parameters, stimulation at no individual channel evoked movements. Each stable stimulation pattern (e.g., blue pulse pattern in Figure 3A, top row) can be envisioned as a *n x m* matrix, when *n* is the contact number and *m* is stimulation duration. The matrix contains ones and zeros, identifying the time of pulse delivered to each contact. To create additional realizations of the stable stimulation patterns (e.g., blue pulse patterns in Supplementary Fig. 5B & C), the assignment of specific peak frequencies to individual channels was randomly shuffled across trials.

For each stable stimulation pattern, we created a corresponding unstable pattern/pair by jittering stimulation pulses within a 20 ms window for each channel and randomly shuffling pulses across channels (e.g., red pulse patterns in Figure 3A, top row and Supplementary Fig. 5B & C). This step created instability in the population pulse pattern by destroying the relative scaling of pulse rates across neurons while also ensuring that both total pulse counts per channel and the mean instantaneous pulse rates across channels were preserved between the stable and unstable patterns. Thus, all subsequent analyses were performed on pairs of trials with stable and unstable stimulation patterns matched in these aspects. We also ensured that the inter-pulse interval was never less than 2 ms (i.e., the peak frequency never exceeded 500 Hz) for any stimulation train. The stimulation patterns were generated offline and the pattern trains corresponding to each trial were introduced 100 ms into the gap period in a gap task. Stable-unstable pairs were randomly interleaved within a block in which roughly 80% of all trials were stimulation trials.

We used the latency of the first saccade after stimulation onset but before stimulation offset as an indicator of the occurrence of a stimulation-evoked saccade (Figure 3B); such saccades exhibited latencies <160 ms. If the microstimulation was ineffective at evoking a saccade, the first movement was typically directed to the target presented after the gap period ended; in such cases, the saccade was produced >400 ms after stimulation onset. We estimated the relative likelihood of evoking a movement by examining trials in which only one of the stable or unstable pulse patterns in a pair yielded a stimulation-evoked saccade (also see Figure 3C and next section). This was quantified based on the number of trials pairs in which only the stable pattern evoked a movement during the window of stimulation (= N_s+us−_, number of points in the blue shaded region in Figure 3B) and the number of trial pairs in which only the unstable pattern evoked a movement (= N_us+s−_, number of points in the red shaded region in Figure 3B). The relative likelihood of evoking a movement with the stable pattern was defined as

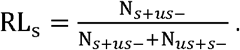

RL_S_ ranges from 0 to 1 and is symmetric with a neutral value of 0.5. To estimate the significance of this estimate, we performed permutation tests with respect to a surrogate dataset in which trials were randomly assigned stable/unstable labels. This lowered the expected estimate of relative likelihood to purely chance levels (Figure 3C).

### Discriminability of population patterns - neural activity and microstimulation

To determine whether the visual and premotor bursts are discriminable using static, non-temporal features of population activity, we performed factor analysis (FA) on 50 ms snippets of activity from these two epochs (visual epoch: 50 ms centered around peak response, motor epoch: 50 ms before saccade onset). We used DataHigh(30), a dimensionality reduction and visualization toolbox written in Matlab, to perform FA on the 16- or 24-dimensional laminar recordings and visualize the projections onto the top latent dimensions. The top 3 dimensions accounted for >95% of the variance in neural states for all our datasets (eigenspectrum not shown), and a 3-dimensional projection is shown for an example dataset in Figure 4A, with the visual and premotor snippets coded in different colors. We also performed FA on the microstimulation patterns (the “states” were computed across the entire 150-ms stimulation duration). Almost all dimensions were needed to account for >95% of the variance in the stimulation patterns, owing to the randomized assignment of frequencies across contacts on different trials. Thus, we chose an arbitrary 3-dimensional projection for visualization in Figure 4B; however, note that the qualitative result did not depend on the projection. Since we used this step purely for visualization, we did not perform any subsequent analysis on the estimated FA dimensions.

Next, we used a linear discriminant analysis (Fisher’s LDA)(*60*) to assess a decoder’s ability to discern whether its inputs signify a movement command at the population level, based on static population features alone, and how this discriminability relates to temporal stability. We first trained a binary LDA classifier on the visual and premotor 50 ms snippets described above in the native neural space. LDA finds a projection defined by a hyperplane that maximizes the separation between the two classes:

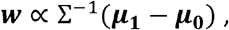

where 0 and 1 are the two class labels, the vector ***w*** is normal to the hyperplane defining the class boundary, 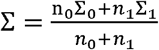, the pooled covariance matrix derived from the within-class covariance matrices Σ_o_ and Σ_1_,n_0_ and n_1_ are class occupancies, and ***μ***_**0**_ and ***μ***_**1**_ the class means. We subjected the linear scores

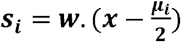

obtained from the LDA projections to a softmax transformation to compute class probabilities:

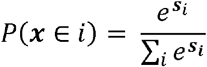

In all cases, we used *k*-fold crossvalidation to test LDA performance (on the classification of visual and premotor population states, and on stimulation patterns described below), with the choice of *k* dictated by the total number of trials in the dataset such that each fold (test set) had at least 10 trials. We quantified classifier performance using the Matthews Correlation Coefficient (MCC)(*61*), since accuracy can be significantly biased and have high variance for classification problems with unequal number of training or test points(*62*), which was the case with the stimulation patterns below. Instead, MCC takes the complete confusion matrix into account to quantify binary classifier performance as:

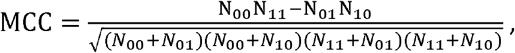

where N_ij_ represents the number of instances where a trial in class *i* is classified to be in class *j* in the classification confusion matrix. Similar to a standard correlation coefficient, MCC values range from -1 to 1 (−1, 0, and 1 respectively indicate poor, average, and perfect classification).

To evaluate the discriminability of microstimulation patterns, we split each dataset in 2 different ways. The first split was based on the predefined stable-unstable pattern pairs and was used to classify pulse patterns as stable or unstable based on their population states alone (inset in Figure 4B). The second split was used to classify stimulation patterns as movement-evoking or non-evoking, in order to identify other features of the stimulation patterns that may have impacted movement occurrence. For this analysis, we focused on stable-unstable trial pairs where only one of the stable-unstable pair evoked a movement (Figure 4C). The rationale was that a classifier trained on this subset may allow for the identification of temporal stability as a predictor because of the differential (i.e., not one-to-one) relationship between stability and movement occurrence. We further evaluated whether a dynamic feature of population activity such as temporal stability can be exploited for this classification by retraining the classifier with explicit addition of temporal stability as an input dimension and comparing it with the performance of one without this addition.

We also performed model selection analysis using nested logistic regression models to identify the predictor variable(s) in the microstimulation patterns that best explained the observed behavior. Logistic regression models the log odds of an outcome (i.e., movement versus no movement) as a linear function of desired predictors (population pulse states and temporal stability) as

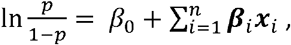

where *p* is the probability of a movement occurring, i.e., P(stimulation evokes movement), ***x*** are the predictors, and ***β*** are the model coefficients. We used a generalized linear model with a logit link function and obtained the maximum likelihood estimate of the coefficients ***β***. We did this for models with one or more predictors chosen from the set of population pulse counts and temporal stability. Since the number of samples and/or predictors can have an effect on the log-likelihood of a given model, we used the Bayesian information criterion (BIC) to estimate model performance, which normalizes log-likelihood by a factor related to the number of samples and predictors. This analysis is shown in Figure 4F.

To explain the variability in behavioral effects of stable/unstable microstimulation patterns across sessions, we also considered whether certain sessions had idiosyncratic patterns of activation across channels for trial subsets in which a movement was evoked. That is, we explored whether differences in pulse rates on subsets of trials could explain the across-sessions range of movement likelihood effects seen in Figure 3C. The rationale was that over many trials, individual channel pulse rates should average out to a narrow distribution of rates given the random uniform sampling on individual trials. Thus, discrepancies from this expected narrow distribution may reflect the effect of pulse rates alone for trial subsets parsed by behavioral effect. We computed relative dispersion for a particular subset of trials as the difference between the dispersion (i.e., pulse count variance across the population) for that subset and dispersion on all trials for that session. That is, *RD*_*subset*_ *= var(x*_*subset*_*) — var(x*_*all*_), where *x* is the distribution across channels of trial-averaged pulse counts for a set of trials, and subset refers to S+US- or US+S- trials. The relative dispersion index was then defined as the relative dispersion for S+US-trials divided by the sum of the two RDs:

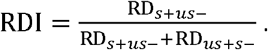

Thus, on sessions where the pulse rates for S+US-trials were closer to the distribution of rates on all trials (closer to the intended uniform sampling of pulse rates across channels), more so than the pulse rates for US+S-trials, the RDI is lower than 0.5 and vice versa. This is an indicator of pulse rate bias in one set of trials versus the other, with the unbiased subset (one with lower RDI) more likely to minimize the influence of pulse rates on the behavioral outcome. This analysis is shown in Supplementary Fig.5.

### Correlation analysis

In order to compare against pooled accumulator mechanisms(*9*), we estimated the correlation structure in our population of neurons and microstimulation patterns. We first computed the spike count correlation across trials for every pair of neurons (or multi-unit clusters, n = 1963 pairs) during the visual and motor epochs as defined above. For the microstimulation patterns, we computed the pulse count correlations for every pair of stimulation channels (n = 899 pairs) during the stimulation window for the stable and unstable patterns separately. Note that in this case, pulse count correlation is equivalent to computing the correlation between accumulation rates (and therefore directly comparable to the accumulator mechanism mentioned above), since the duration of stimulation was fixed and the baseline and peak rates were matched between the stable and unstable conditions.

### Spiking neuron model

We developed a biophysically realistic computational model with spiking neurons to discriminate and read out temporal structure in population activity. The core component of the model was a recurrently connected module comprising a decoder element and a disinhibitor element, putatively representing pontine high-frequency burst neurons and OPNs, respectively(*37*). The decoder and disinhibitor were both modelled as leaky integrate-and-fire (LIF) spiking neurons:

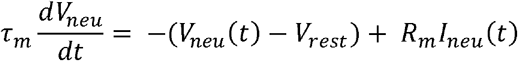

where *V*_*neu*_(*t*) is the membrane potential of the neuron, *V*_*rest*_ is the resting potential, *τ*_*m*_ is the membrane time constant, *R*_*m*_ is the membrane resistance, *I*_*neu*_(*t*) is the net time-varying input to the neuron, and *neu* represents the decoder *(dec)* or disinhibitor *(dis)*. The spiking followed standard threshold crossing rules, i.e., if *V*_*neu*_*(t = t*^*k*^*) > V*_*th*_, where *V*_*th*_ represents the action potential threshold, the membrane potential was reset to *V*_*neu*_*(t = t*^*k*^*) = V*_*reset*_, and the neuron’s spike times [*t*^*i*^] were updated with the *k*^*th*^ spike time *t*^*k*^.

The current input to both the decoder and disinhibitor was composed of excitatory and inhibitory components:

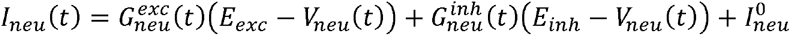

where 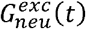 and 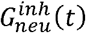 are the respective net excitatory and inhibitory post-synaptic conductances, and *E*_*exc*_ and *E*_*inh*_ are the excitatory and inhibitory reversal potentials representing the contribution of AMPA and GABA channels, respectively. 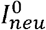 is a constant current term that was set to zero for the decoder and to a fixed value of 4 for the disinhibitor in order to simulate tonic firing without external input.

The decoder received excitatory inputs from the SC in the form of stimulation pulse trains, and inhibitory input from the disinhibitor. The disinhibitor received both excitatory and inhibitory inputs from the SC(*33, 34*) and reciprocal inhibition from the decoder. The net time-varying conductances were modelled as follows:

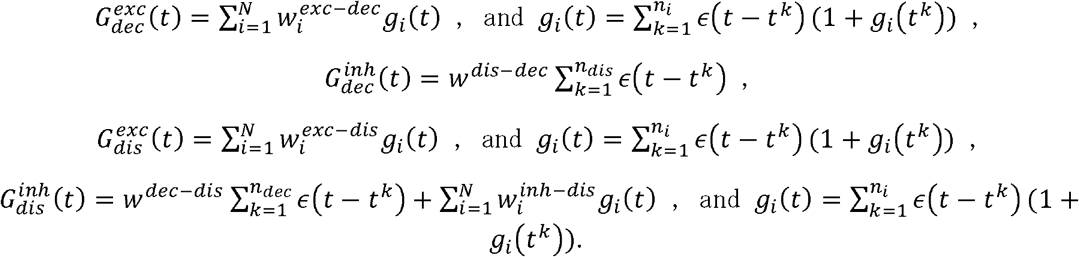

In each equation they appear, *N* is the number of input units (i.e., number of channels with pulse trains), 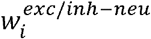 is the excitatory/inhibitory weight from the *i*^*th*^ input unit to the spiking neuron, *n*_*i*_ is the number of pulses in the *i*^*th*^ input unit, *w*^*dl-sdec*^ is the weight from the disinhibitor to the decoder, *w*^*dec-dis*^ is wejght from the decoder to the disinhibitor, an *k*^*th*^ is the time of the *k*^*th*^ input pulse or output spike (depending on which summation term it appears in). The *g*_*i*_ term, when it appears in the sum that produces *g*_*i*_ (as in all but the second conductance equation), represents the state-dependent influence of each incoming pulse input. *∈*(t) is the post-synaptic conductance kernel defined as:

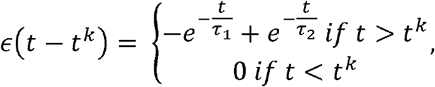

The conductance and input current terms that were dependent solely on the pulse input patterns were pre-computed and fed into a Eulerian solver (time step = 0.1 ms) for the differential equations governing the membrane potential of the spiking decoder and disinhibitor units. Supplementary Table 1 shows the values of the parameters and constants used for the model simulations.

## Acknowledgements

We thank Drs. A. Batista, C. Olson, M. Smith, and B. Yu for scientific discussions and critical feedback on previous versions of the manuscript and J. McFerron for programming assistance. The study was funded by the following NIH R01 grants: EY022854 and EY024831 awarded to N.J.G.

## Funding

The study was funded by the following NIH R01 grants: EY022854 and EY024831.

## Author Contributions

U.K.J and N.J.G conceived and designed the study. U.K.J. performed the experiments, analyzed the data, and performed model simulations. U.K.J and N.J.G wrote the manuscript.

## Competing Interests

The authors declare no competing financial interests.

## Data and code availability

The data that support the findings of this study are available from the corresponding author(s) upon reasonable request. The custom MATLAB code is available from the authors upon reasonable request.

**Supplementary Figure 1.**
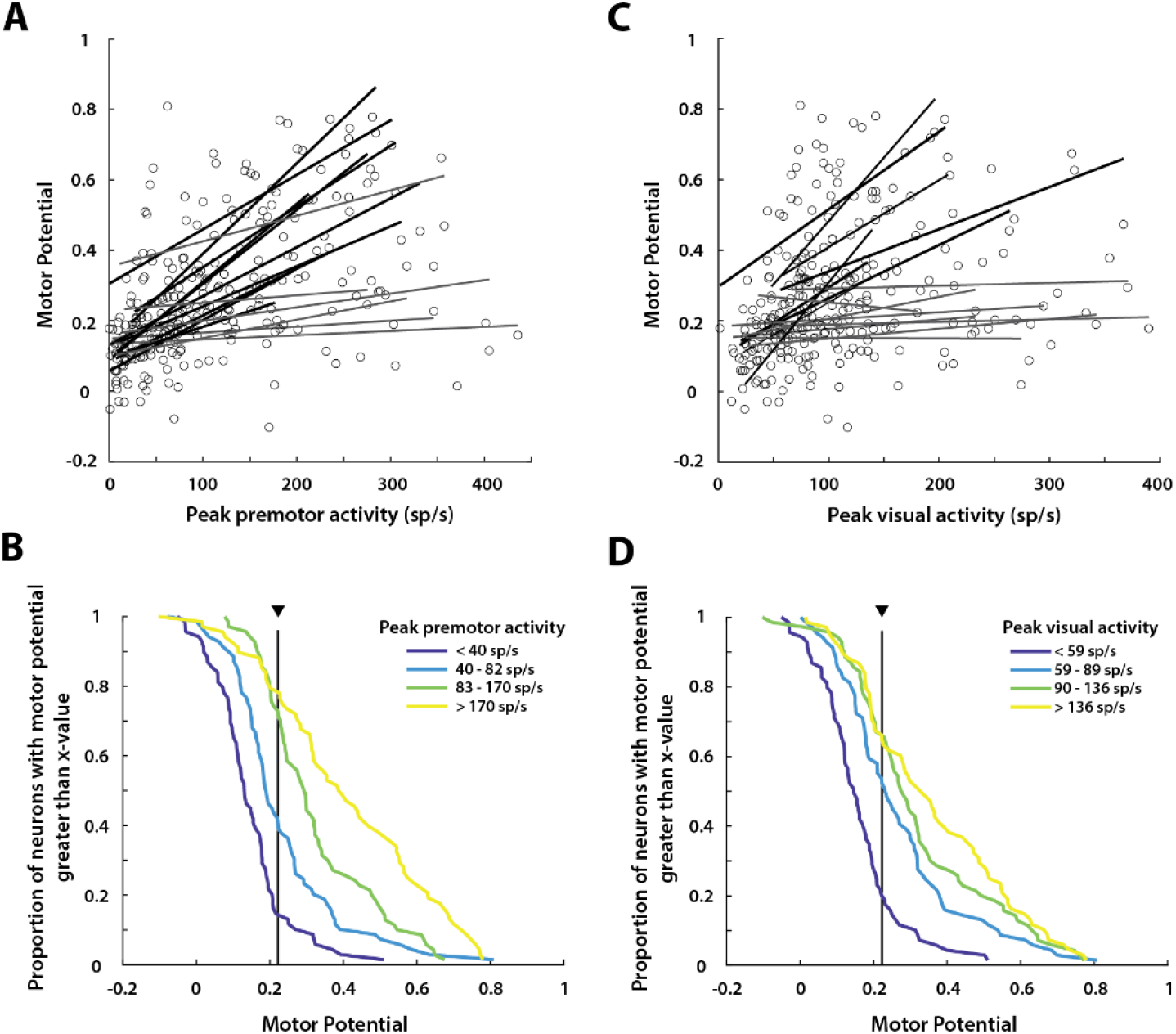
Strong saccade drivers also have high visual activity. **A**. Scatter plot of the motor potential, calculated as the peak correlation between saccade velocity and preceding premotor activity (see Methods), as a function of peak premotor burst for individual neurons. Each point represents a neuron/MUA recorded on an individual channel, and each line is the linear regression fit for the points from an individual, laminar probe recording session (bold lines indicate significant trends, p < 0.01). **B**. Neurometric plot of the proportion of neurons in each of 4 quartiles split based on peak premotor activity that have a motor potential (as defined above) higher than the abscissa. The traces are colored based on their quartile as indicated in the legend. The vertical line and arrowhead indicate the median motor potential. For example, roughly 80% of the neurons in the highest premotor activity quartile (yellow trace) show an activity-velocity correlation above the median and are thus considered strong saccade drivers. **C**. Scatter plot similar to **A** except the abscissa shows peak visual activity. A strong correlation between the neurons’ peak visual activity and its motor potential, which is still computed using its premotor activity, is observed for several sessions and in the overall distribution. **D**. Neurometric plot similar to **B** of the proportion of neurons in each of 4 quartiles split based on peak *visual* activity that have a motor potential higher than the abscissa. The majority of neurons with high visual activity (>60% in top 2 quartiles, green and yellow traces) exhibit motor potential above the median (vertical line and arrowhead), indicating that neurons with high visual activity also strongly influence saccade kinematics and thus are candidates for causal drivers of the movement.

**Supplementary Figure 2.**
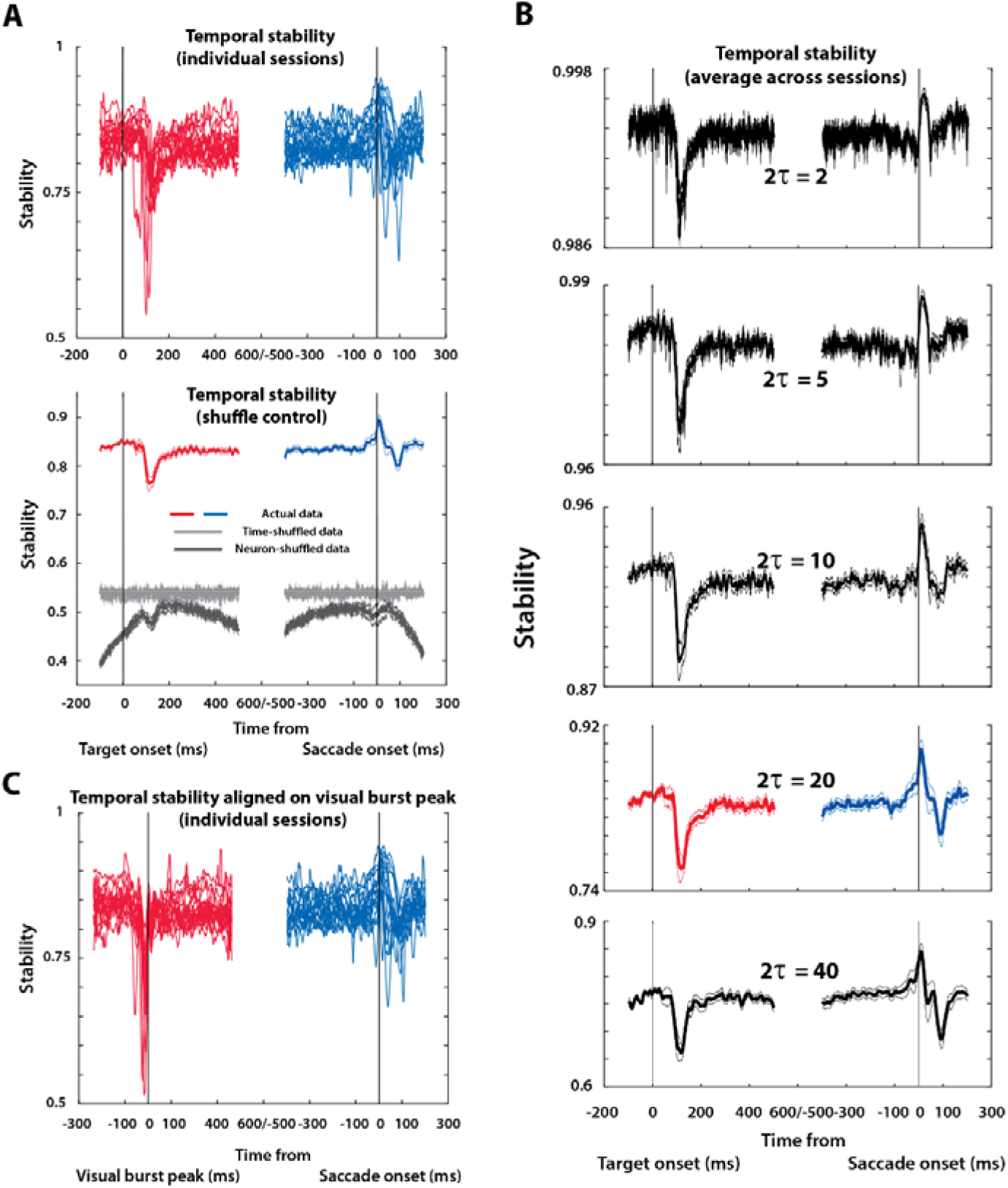
Temporal stability profiles within and across laminar recording sessions. **A**. Top row: Trial-averaged temporal stability traces for individual sessions (thin traces) have different baselines, possibly due to varying noise levels, spike isolation differences, etc. The baselines were shifted to the across-session mean level before session-averaging in Figure 1D. Note that the raw and not baseline-shifted values were used for statistical analyses. Bottom row: Temporal stability averaged across trials and sessions for two types of shuffled datasets. The light gray traces (mean +/- s.e.m.) are for time-shuffled data where relative activity levels between neurons are preserved but time points during the trial are randomly re-ordered. The dark gray traces are for neuron-shuffled data where the instantaneous population firing rate is preserved but the firing rate distribution is shuffled across neurons at each time point. The colored traces show the session-averaged temporal stability (as in Figure 1D) for comparison. **B**. In order to ensure that the choice of *τ* did not have an undue effect on our results, we computed stability for several values of *τ*. The panels show temporal stability profiles for (from top to bottom), *τ* = 1, 2.5, 5, 10, and 20, respectively. The colored traces for *τ* =10 are the ones shown in the Figure 1D. As evident, the absolute magnitude of stability was inversely related to *τ*, a consequence of the smooth and continuous nature of the trajectory. In other words, since the state of the neural population evolves smoothly, it must traverse intermediate states in order to move from one state to another, resulting in greater similarity between state vectors close together in time than those further apart. However, the relative shape of stability profile was largely preserved for *τ* > 5. **C**. Similar to top row of panel A but for temporal stability aligned to peak of visual burst.

**Supplementary Figure 3.**
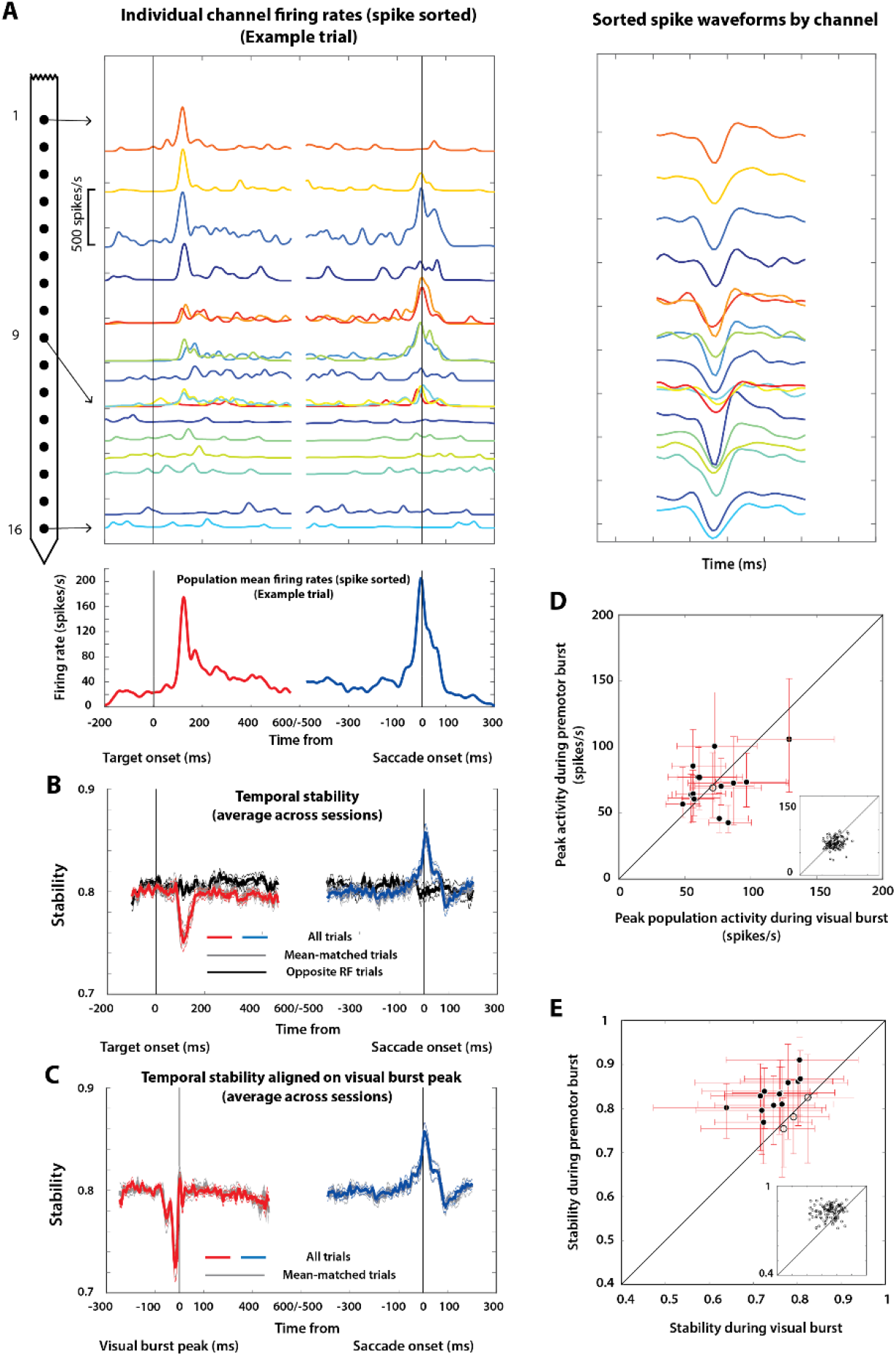
Spike sorting does not affect temporal structure findings. **A**. Similar to Figure 1B but for spike sorted data. Top left: Example laminar recording trial showing activity from 16 channels aligned on target (left column) and saccade (right column) onsets. Multiple spike sorted units in a channel (if any) are shown as different traces. Bottom: Population activity averaged across the 16 channels for the example trial. Top right: Corresponding mean waveforms of spike sorted units in each channel, with multiple units from a channel shown as different traces. **B**. Temporal stability for spike sorted laminar recordings shows a similar trend to the unsorted multi-unit activity data in the main figures (dip during the visual epoch and increase in the premotor epoch, p < 0.01 for Wilcoxon signed-rank test with respect to baseline, cf. Figure 1D). **C**. Temporal stability for spike sorted data aligned to peak visual burst, similar to Figure 1E. **D**. Scatter plot of mean peak population firing rate from spike sorted data for individual sessions, with trials from an example session shown in the inset (cf. Figure 1G). **E**. Similar to D but for temporal stability in a 20 ms window preceding the peak visual and premotor bursts (cf. Figure 1F). In D and E, filled circles indicate significant difference between the corresponding values computed during the visual burst and the premotor burst (p < 0.01, Wilcoxon signed-rank test).

**Supplementary Figure 4.**
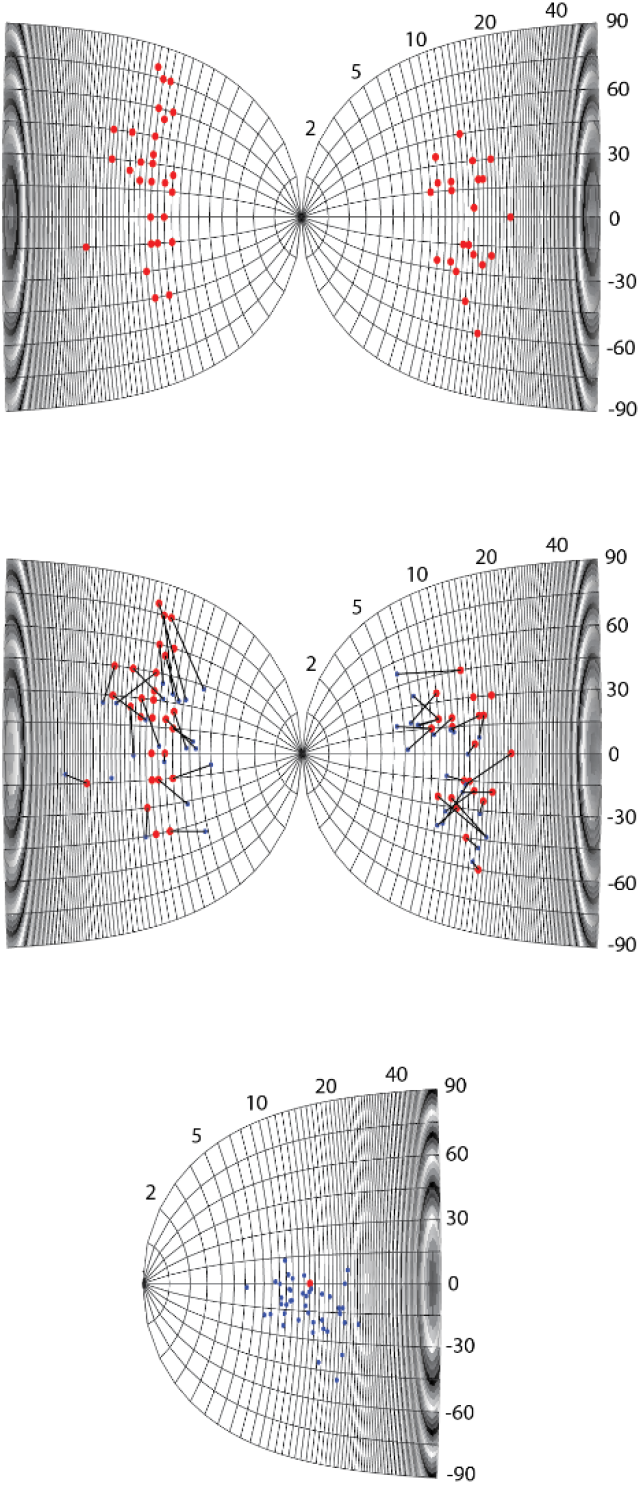
Reconstructed locations of neurons in pseudo-population on SC map. *Top row -* Target loci (red points) mapped from visual space to SC coordinates across all experimental sessions. The locations on the SC were computed using traditionally established transformations between visual space and SC tissue coordinates (see Methods). *Middle row -* Endpoints of the stimulation-evoked saccade vector at the recording site (blue points) shown with the target loci from above. The stimulation vector provides a proxy for the RF center of the recorded neuron. Neuron-target pairs (blue-red) from individual sessions are connected using black lines (unconnected points did not have the corresponding stimulation/target data). *Bottom row -* The active pseudo-population on the SC map. The red locations from the previous panels were referenced to one arbitrarily selected location on the SC map (here, R=15, theta = 0) and the blue locations were translated accordingly. Points from both colliculi are shown on a single SC for the sake of clarity.

**Supplementary Figure 5.**
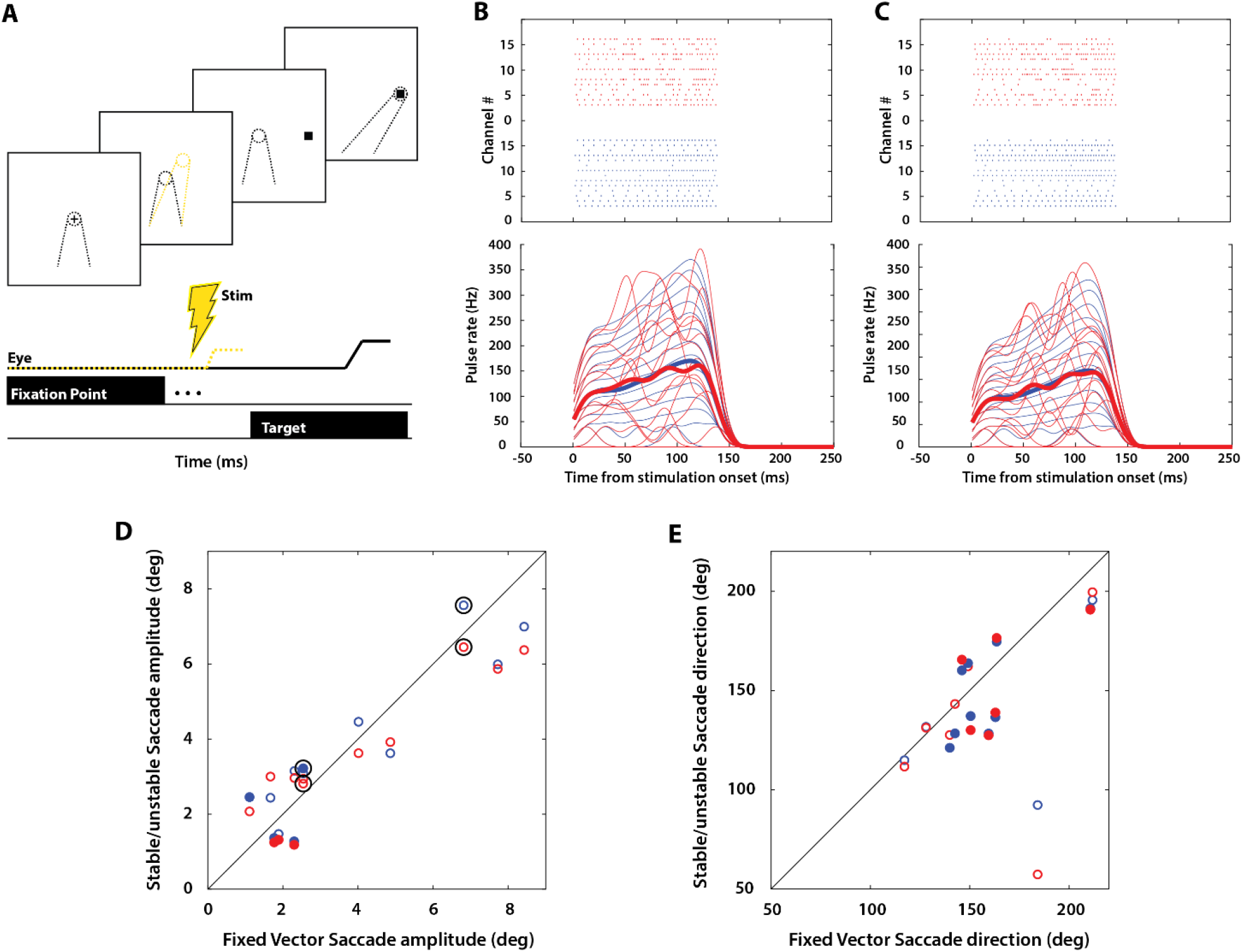
Microstimulation experiment - task, examples, and evoked saccade metrics. **A**. Schematic of the gap task used in the patterned microstimulation experiments. **B** and **C**. Two more examples of pulse rasters and pulse rates for individual channels for stable (blue) and unstable (red) pairs of stimulation patterns (cf. Figure 3A). **D**. Mean stimulation-evoked saccade amplitude for stable (blue points) and unstable (red points) stimulation patterns plotted against the fixed vector saccade amplitude obtained from near-threshold constant frequency stimulation (see Methods). The fixed-vector saccade is also known as the site-specific saccade. Each point represents one session for that condition. The diagonal represents the unity relationship. Filled circles denote a significant difference of the stable/unstable saccade amplitude from the fixed vector saccade (p < 0.01, Wilcoxon rank-sum test). Points circumscribed by black circles denote sessions where the saccade amplitudes on stable and unstable trials were significantly different from each other (p < 0.01, Wilcoxon signed-rank test). **E**. Similar to D (including criteria for statistical significance), but for stimulation-evoked saccade direction.

**Supplementary Figure 6.**
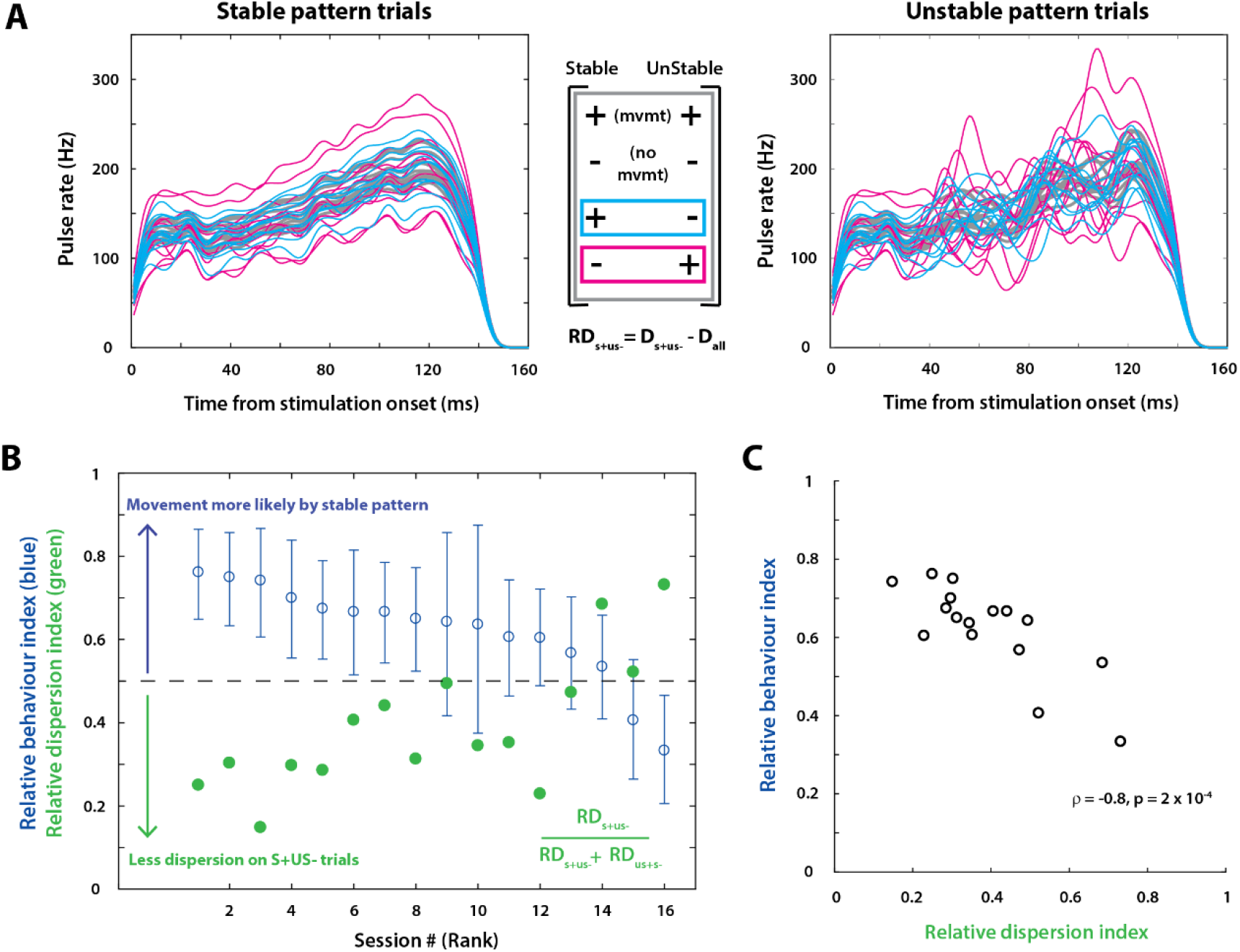
Variability in behavior is partly explained by variability in stimulation pulse rates across subsets of trials. **A**. Individual channel pulse rates averaged across specific subsets of trials for an example session. *Left -* Average stable pattern for the set of trials in which, only the stable pattern evoked a movement out of the stable-unstable pair (S+US- pairs, cyan traces), only the unstable pattern evoked a movement (S-US+ pairs, magenta traces), and all trials (gray traces). *Middle -* Schematic of which trial pairs are plotted in the left and right columns, highlighted in the appropriate color. Relative dispersion is calculated as the across-channel dispersion (variance) of the pulse counts for a particular subset (e.g., S+US-) minus the dispersion of the pulse counts for all trials. *Right* – Average unstable pattern for the three subsets of trials as in the left column. **B**. Relative dispersion index computed as in the formula in the inset, plotted for each session (green points). Values *less than* 0.5 indicate that there was less dispersion in the S+US-trials, and trials with lower dispersion are less dependent on pulse rate effects. The behavioral effect (from Figure 3C) is overlaid for comparison (blue points and error bars). The sessions are ordered by behavioral effect size as in Figure 3C. **C**. Scatter plot showing correlation between behavioral effect size and relative dispersion index. The sessions with negative (ordinate < 0.5) or non-existent effect sizes in behavior are also the ones with the highest dispersion index, indicating that those sessions were dominated by pulse rate effects and less likely to reveal an effect of temporal structure.

**Supplementary Figure 7.**
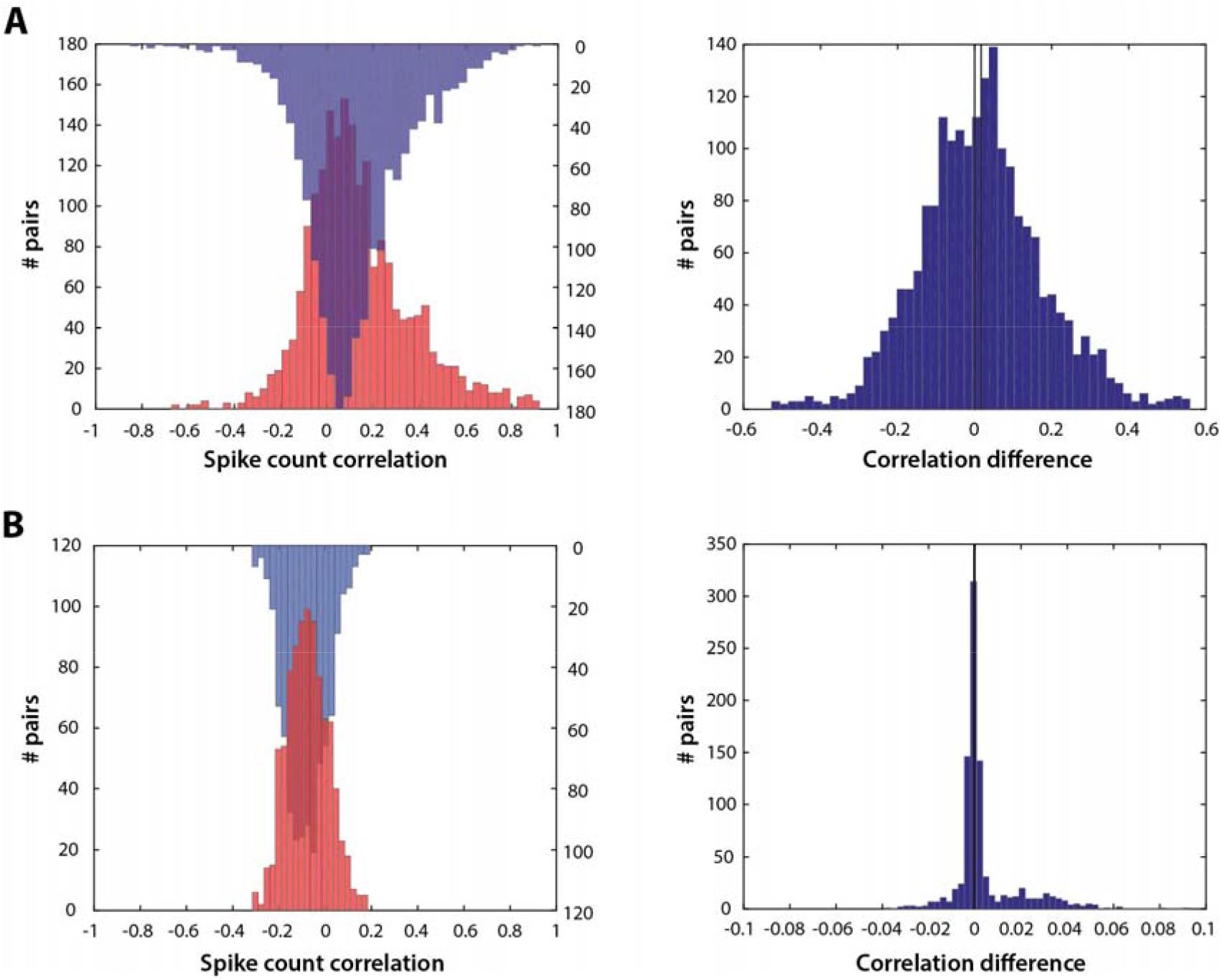
Across-trial spike and pulse count correlations. **A**. Left: Histograms of spike count correlations for all pairs of channels (n = 1963) for the visual and premotor epochs (red and blue histograms, respectively). One of the histograms is shown inverted for the sake of clarity. Right: Distribution of differences between visual and premotor epoch correlations computed for each pair. Positive values indicate larger spike count correlations for spikes in the visual epoch. The two vertical lines represent zero and the mean of the distribution (left and right lines, respectively). **B**. Same as A for pulse counts correlations for all pairs of stimulation channels (n = 899) for the stable (blue) and unstable (red) patterns, with the corresponding difference histogram shown on the right. Positive values indicate higher pulse count correlations in the unstable stimulation pattern.

**Supplementary Figure 8.**
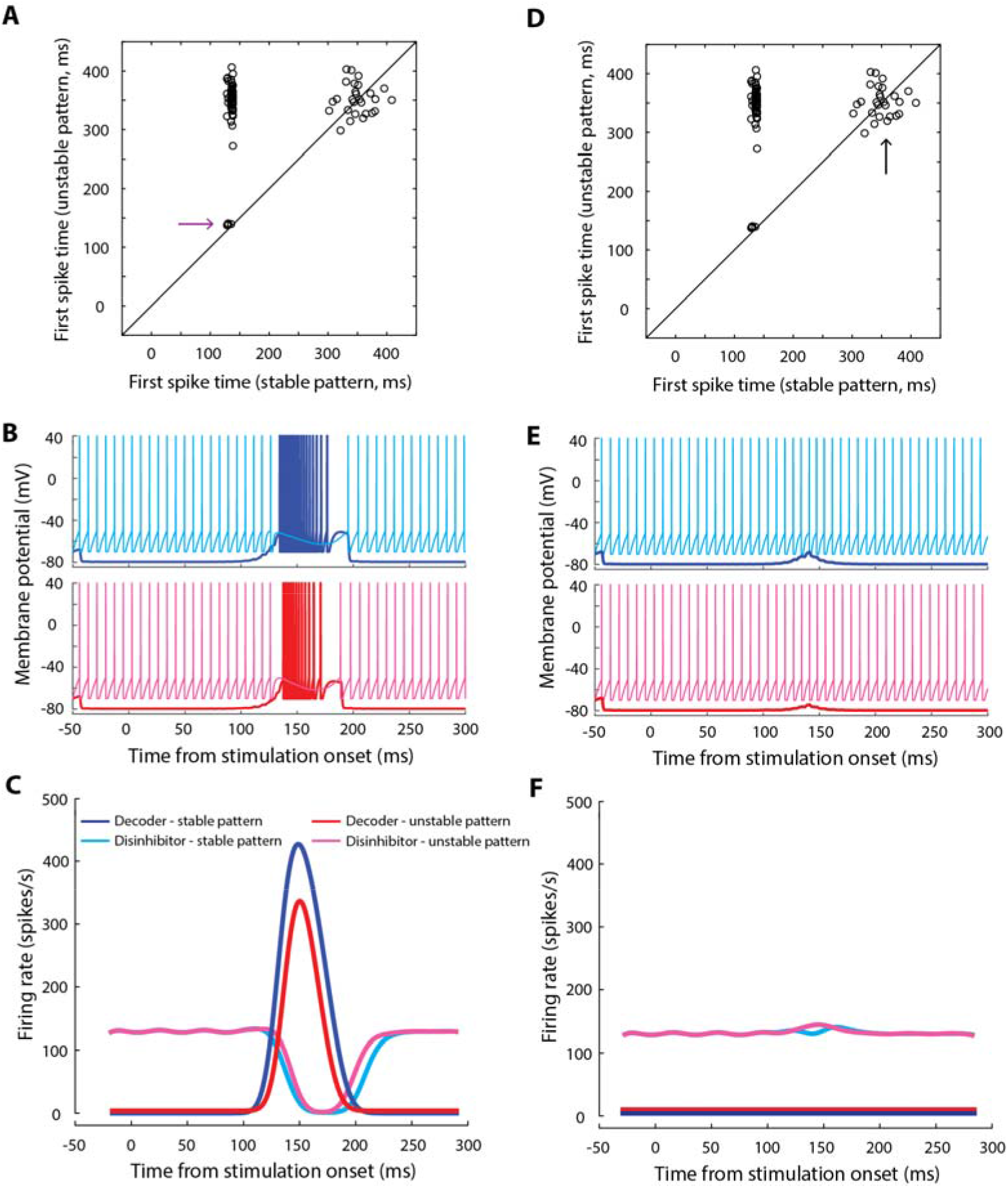
Performance of spiking neuron model for various simulation outcomes with stable and unstable pulse patterns. **A**. Scatter plot of first spike latencies in the decoder (putatively representing saccade initiation) for stable versus unstable inputs from matched pairs (same as Figure 5C). The next two panels are based on the trial pairs highlighted by the arrow (in which both stable and unstable patterns produced a saccade). **B**. Simulated membrane potential of the decoder (blue and red traces) and disinhibitor (cyan and magenta traces) for an example matched trial pair from the subset in panel A, for stable (top row) and unstable (bottom row) input patterns. **E**. Average activity of the disinhibitor (cyan and magenta traces, stable and unstable inputs, respectively) and decoder (blue and red traces, stable and unstable inputs, respectively) for all trial pairs in which both stable and unstable input patterns produced a movement. **D**. Same as panel A, except for the focus on trial pairs (arrow). The next two panels are based on the trial pairs in which neither stable nor unstable patterns produced a saccade. **E**. Simulated membrane potential of the decoder and disinhibitor for an example matched trial pair from the subset in panel D. Layout and color scheme as in panel B. **F**. Average activity of the disinhibitor and decoder for all trial pairs in which neither stable nor unstable input patterns produced a movement. Color scheme as in panel C. Note the small increase in disinhibitor activity around the time when a movement would have normally occurred, reminiscent of the slight increase in OPN activity during visual input(*38*).

**Supplementary Figure 9.**
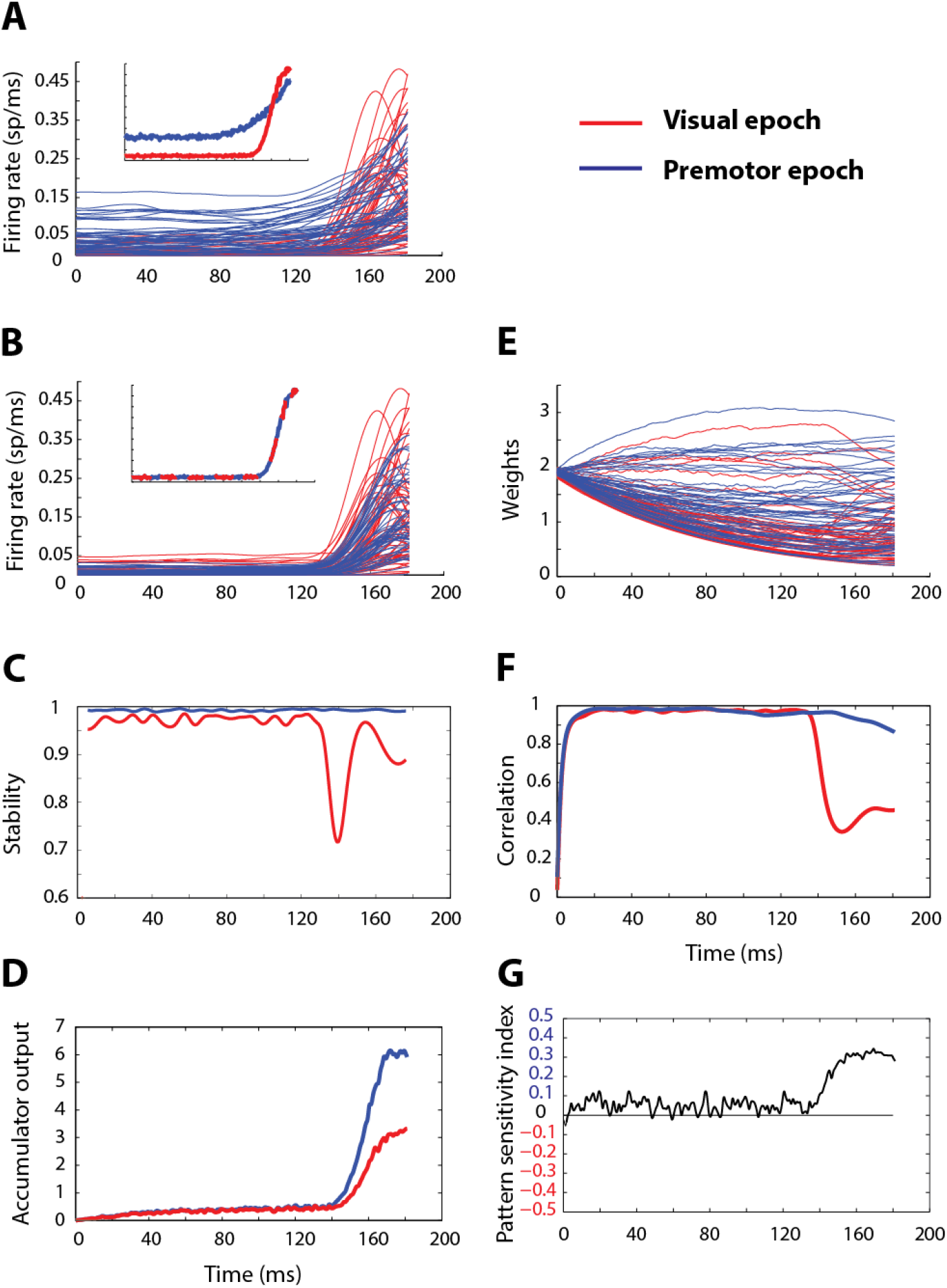
Leaky accumulator with facilitation (LAF) model can discriminate population temporal structure. **A**. Inputs to the model from SC visuomovement neurons. Raw unstable (red) and stable (blue) input profiles (inset – population means). The two sets of inputs are 180 ms snippets taken from the visual and premotor bursts, respectively, in the spike density profiles shown in Figure 2A. **B**. Mean-matched input profiles and population means. **C**. Temporal stability of the two mean-matched populations. **D**. Output of the LAF accumulator in response to the stable and unstable model inputs. **E**. Evolution of synaptic weights for the two conditions. **F**. Correlation between instantaneous weights and input rates for the two conditions. **G**. Pattern sensitivity index of the model’s ability to discriminate between the two types of inputs. Values in the top half of the plot indicate higher sensitivity (faster accumulation) to stable population input.

**Supplementary Table 1.** Values of the constants and parameters used for the spiking neuron model.

| Biophysical constants | Values |
| --- | --- |
| $\tau_m$ | 10 ms |
| $R_m$ | 10 Mohms |
| $V_{th}$ | -50 mV |
| $V_{rest}$ | -65 mV |
| $V_{reset}$ | -70 mV |
| $E_{exc}$ | 50 mV |
| $E_{inh}$ | -80 mV |
| Model parameters |  |
| $\tau_1$ | 2 ms |
| $\tau_2$ | 10 ms |
| $w^{exc-dec}$ | $5 \times 10^{-6} \text{ uniform}[0,2]$ |
| $w^{dis-dec}$ | 5 |
| $w^{exc-dis}$ | $0.2 w^{exc-dec}$ |
| $w^{inh-dis}$ | $0.8 w^{exc-dec}$ |
| $w^{dec-dis}$ | 0.1 |
| Numerical parameters |  |
| $dt$ (time step) | 0.1 |

## S1. Inferring population dynamics from single-unit recordings

The main text and methods describe how we reconstructed a pseudo-population from unit recordings. Using knowledge of the RF center of a given session’s local population obtained from microstimulation, and known transformation from visual space to SC tissue coordinates, here we reconstruct the pseudo-population on the SC map.

We first transferred target locations (*R*_*T*_, *θ*_*T*_) inside the RF for each neuron onto the SC map using the following transformations(*59*) -

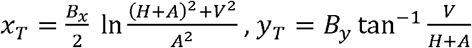, where,

*H* = *R*_*T*_ COS *θ*_*T*_, = *R*_*T*_ sin *θ*_*T*_, and *A =* 3, *B*_*x*_ *=* 1.4, *B*_*y*_ *=* 1.8.

The transformed locations *(x*_*T*_,*y*_*T*_) on the bilateral SC map are shown in Supplementary Fig. 3A, top row. In order to move these target locations to one pseudo-population, we need to identify where these points reside in the RF of each neuron or local population. We used the endpoints of the site-specific microstimulation-evoked saccades as a proxy for the respective RF centers. The transformed endpoints, with their locations relative to the corresponding target locations, are shown in Supplementary Fig. 3A, middle row. We picked an arbitrary location in the visual field as the RF center of the pseudopopulation, (*R*_*c*_, *θ*_*c*_) = (*15,0*). We used the fact that the size of the active SC population is invariant regardless of the encoded saccade to preserve relative distances in tissue space between a site’s target location and RF centre, and translated the transformed microstimulation endpoints to the common pseudo-population center, along with the respective target locations. To construct one active population from sites gleaned from both hemifields (and therefore both colliculi), we reflected the coordinates from one colliculus onto the other. The resulting pseudo-population (Supplementary Fig. 3A, bottom row) shows a fairly representative sampling of neurons from the pseudo-population, with its extent consistent with a large (25% of SC tissue) active population, albeit one that is biased to one side of the population.

## S2. Leaky accumulator with facilitation (LAF) model

We developed an alternative computational model to demonstrate how the temporal structure of population activity could be used by neural circuits to gate decisions such as movement initiation. Similar to the spiking neuron model presented in the main text, a core principle behind this model is the hypothesis that signal integration is stronger when the temporal structure in input activity is stable, which allows the decoder neuron to reach threshold during the high frequency burst that triggers the movement. We constructed an accumulator as an abstraction of a neuron receiving population inputs through its network of dendrites. The total synaptic current *I*(*t*) and the firing rate *v*(*t*) of the accumulator were defined as

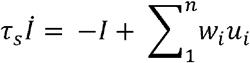

and

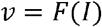

where *u*_*i*_ and *W*_*i*_ are the instantaneous firing rate and synaptic gain of the *i*^*th*^ input neuron, and *τ*_*s*_ is the time constant of the synaptic current. The output firing rate of the single decoder neuron *v*(*t*) can be described by a standard monotonic function (e.g., linear or sigmoid) applied to the net current(*63*). The family of stochastic, leaky accumulator models that integrate the firing rate of neurons toward a threshold criterion has been commonly used to explain reaction times, decision making, and perception(*9,64-66*). We used the following heuristic in order to extend this framework to incorporate temporal structure. In order to assess stability, the decoder neuron must keep track of the short term history of the population activity, use this “memory” to evaluate stability, and respond selectively when the activity pattern is deemed stable over some time scale. We implemented these requirements by introducing short-term plasticity in the form of facilitatory connections(*67, 68*) from the input population onto the output unit (accumulator). The change in connection strength or gain of each neuron on the decoder neuron *w*_*t*_ can be defined by a differential equation that incorporates a Hebbian-like learning rule and leak current:

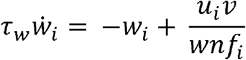

The Hebbian-learning component, *u*_*i*_*v* describes the change in weight coupled to the firing rates of the *i*^*th*^ pre-synaptic neuron and the post-synaptic accumulator neuron. *τ*_*W*_ is the time constant of the weight parameter. Since this version of the model contains only excitatory connections, the weight parameter can exhibit unbounded accumulation, which can be controlled by incorporating normalization. We normalized the Hebbian-learning component *u*_*i*_ *v* with the contribution of that neuron to the total resource pool(*69*). The resource pool available for facilitation at any given time was defined as the sum of the Hebbian-learning component across all input units 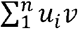. The contribution of a neuron to the output rate *v*(*t*) then determines its contribution to the overall pool, giving rise to the weight normalization factor:

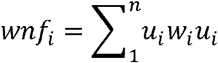

For this model simulation, we used visuomovement neuron activity drawn from the SC pseudopopulation as inputs (Supplementary Fig. 8A). We created two sets of input snippets from the visual and premotor bursts, each 180 ms in length. For the visual burst, we used the activity upto the peak in the population response. For the premotor burst, we used activity upto 35 ms before saccade onset (which was the time when the response magnitude reached the same level as the peak of the visual burst). In order to control for the effect of average firing rate on the accumulation, we mean-matched the input profiles as follows. We divided the activations in the premotor input set at each instant by the mean activation of the visual inputs at the corresponding instant. That is,

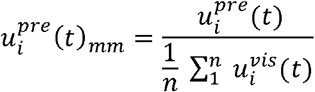

where 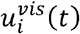 and 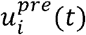 are the activity of neuron *i* at time *t* in the visual and 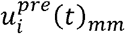 is the premotor input sets, respectively, and is the premotor input instantaneously rescaled to match the mean of the visual input (Supplementary Fig. 8B). This ensured that we isolated the effect of temporal structure (Supplementary Fig. 8C), independent of firing rate, on the model’s response.

The accumulator increased its activity at a faster rate in response to the stable pattern compared to the unstable pattern (Supplementary Fig. 8D), suggesting that the network was able to discriminate between two types of population patterns even though the net input drive was identical. How critical is facilitation to this function? Like the inputs themselves, the weights also showed greater fluctuation during the visual burst (Supplementary Fig. 8E). Consistent with our heuristic, the weights tracked the input rates when the input was stable, but this correlation dropped away when the input was unstable (Supplementary Fig. 8F). Note that the shape of the correlation profile is not unlike the stability profile in Supplementary Fig. 8C. Therefore, facilitation allows the weights to retain the memory of a pattern over the time scale of tens of milliseconds, but the memory can be scrambled by a fluctuating input pattern.

We also quantified the ability of the synaptic weights to track the inputs by computing the Pearson’s correlation between the weight and input vectors at each time point (Supplementary Fig. 8G). To quantify the accumulator’s ability to discriminate temporal pattern in population input, we computed a pattern sensitivity index *(psi)* as

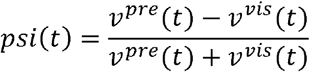

where *v*(*t*) is the output of the accumulation for the corresponding inputs.

The results here suggest that the accumulator is driven more strongly by the stable premotor burst, even when other features of population activity remain the same, providing yet another biophysical mechanism by which temporal structure could be read out.

## Notes

**Conflict of interest:** The authors declare no competing financial interests.

### Competing Interest Statement

The authors have declared no competing interest.

### Summary of Updates

The significant changes from the previous version are as follows: 1) New detailed introduction. 2) Analysis of spike sorted data replicating the main results in Figure 1 (Supplementary Figure 2) 3) Complete revision of Figure 2 with more standard pseudo-population analyses, and express saccade data. 4) Minor updates to the text and other figures.

## References

1. R. H. Wurtz, M. A. Sommer, M. Paré, S. Ferraina, Signal transformations from cerebral cortex to superior colliculus for the generation of saccades. Vision Res 41, 3399–3412 (2001).

2. C. A. Buneo, M. R. Jarvis, A. P. Batista, R. A. Andersen, Direct visuomotor transformations for reaching. Nature 416, 632–636 (2002).

3. V. Gallese, L. Fadiga, L. Fogassi, G. Rizzolatti, Action recognition in the premotor cortex. Brain : a journal of neurology 119 (Pt 2), 593–609 (1996).

4. C. K, Rodgers, D. P. Munoz, S. H. Scott, M. Pare, Discharge properties of monkey tectoreticular neurons. J Neurophysiol 95, 3502–3511 (2006).

5. M. A. Segraves, Activity of monkey frontal eye field neurons projecting to oculomotor regions of the pons. J Neurophysiol 68, 1967–1985 (1992).

6. D. P. Hanes, J. D. Schall, Neural control of voluntary movement initiation. Science 274, 427–430 (1996).

7. R. Ratcliff, J. N. Rouder, Modeling response times for two-choice decisions. Psychol Rev 9, 347–356(1998).

8. J. J. Jantz, M. Watanabe, S. Everling, D. P. Munoz, Threshold mechanism for saccade initiation in the frontal eye field and superior colliculus. J Neurophysiol 109, 2767–2780 (2013).

9. B. Zandbelt, B. A. Purcell, T. J. Palmeri, G. D. Logan, J. D. Schall, Response times from ensembles of accumulators. Proceedings of the National Academy of Sciences of the United States of America 111, 2848–2853 (2014).

10. B. M. Yu et al., Gaussian-process factor analysis for low-dimensional single-trial analysis of neural population activity. J Neurophysiol 102, 614–635 (2009).

11. K. V. Shenoy, M. Sahani, M. M. Churchland, Cortical control of arm movements: a dynamical systems perspective. Annu Rev Neurosci 36, 337–359 (2013).

12. M. M. Churchland et al., Neural population dynamics during reaching. Nature 487, 51–56 (2012).

13. M. T. Kaufman, M. M. Churchland, S. I. Ryu, K. V. Shenoy, Cortical activity in the null space: permitting preparation without movement. Nature neuroscience 17, 440–448 (2014).

14. G. F. Elsayed, A. H. Lara, M. T. Kaufman, M. M. Churchland, J. P. Cunningham, Reorganization between preparatory and movement population responses in motor cortex. Nat Commun 7, 13239 (2016).

15. A. Afshar et al., Single-trial neural correlates of arm movement preparation. Neuron 71, 555–564 (2011).

16. N. Miyashita, O. Hikosaka, Minimal synaptic delay in the saccadic output pathway of the superior colliculus studied in awake monkey. Exp Brain Res 112, 187–196 (1996).

17. U. K. Jagadisan, N. J. Gandhi, Removal of inhibition uncovers latent movement potential during preparation. Elife 6, (2017).

18. I. Smalianchuk, U. K. Jagadisan, N.J. Gandhi, Instantaneous Midbrain Control of Saccade Velocity. J Neurosci 38, 10156–10167 (2018).

19. P. J. May, J. D. Porter, The laminar distribution of macaque tectobulbar and tectospinal neurons. Vis Neurosci 8, 257–276 (1992).

20. T. Branco, B. A. Clark, M. Hausser, Dendritic discrimination of temporal input sequences in cortical neurons. Science 329, 1671–1675 (2010).

21. A. G. Carter, G. J. Soler-Llavina, B. L. Sabatini, Timing and location of synaptic inputs determine modes of subthreshold integration in striatal medium spiny neurons. J Neurosci 27, 8967–8977 (2007).

22. M. A. Harvey, H. P. Saal, J. F. Dammann, 3rd, S. J. Bensmaia, Multiplexing stimulus information through rate and temporal codes in primate somatosensory cortex. PLoS biology 11, el001558 (2013).

23. E. M. Trautmann et al., Accurate Estimation of Neural Population Dynamics without Spike Sorting. Neuron 103, 292–308 e294 (2019).

24. C. J. Bruce, M. E. Goldberg, Primate frontal eye fields. I. Single neurons discharging before saccades. J Neurophysiol 53, 603–635 (1985).

25. G. B. Stanton, M. E. Goldberg, C. J. Bruce, Frontal eye field efferents in the macaque monkey: II. Topography of terminal fields in midbrain and pons. J Comp Neurol 271, 493–506 (1988).

26. D. P. Munoz, R. H. Wurtz, Fixation cells in monkey superior colliculus. I. Characteristics of cell discharge. J Neurophysiol 70, 559–575 (1993).

27. A. K. Moschovakis et al., An anatomical substrate for the spatiotemporal transformation. J Neurosci 18, 10219–10229 (1998).

28. Z. M. Hafed, L. Goffart, R. J. Krauzlis, A neural mechanism for microsaccade generation in the primate superior colliculus. Science 323, 940–943 (2009).

29. D. L. Kimmel, T. Moore, Temporal patterning of saccadic eye movement signals. J Neurosci 27, 7619–7630 (2007).

30. B. R. Cowley et al., DataHigh: graphical user interface for visualizing and interacting with highdimensional neural activity. Journal of neural engineering 10, 066012 (2013).

31. C. Massot, U. K. Jagadisan, N. J. Gandhi, Sensorimotor transformation elicits systematic patterns of activity along the dorsoventral extent of the superior colliculus in the macaque monkey. Commun Biol 2, 287 (2019).

32. R. P. Heitz, J. D. Schall, Neural mechanisms of speed-accuracy tradeoff. Neuron 76, 616–628 (2012).

33. N. J. Gandhi, E. L. Keller, Spatial distribution and discharge characteristics of superior colliculus neurons antidromically activated from the omnipause region in monkey. J Neurophysiol 78, 2221–2225 (1997).

34. K. Yoshida, Y. Iwamoto, S. Chimoto, H. Shimazu, Saccade-related inhibitory input to pontine omnipause neurons: an intracellular study in alert cats. J Neurophysiol 82, 1198–1208 (1999).

35. R. A. Deisz, G. Fortin, W. Zieglgansberger, Voltage dependence of excitatory postsynaptic potentials of rat neocortical neurons. J Neurophysiol 65, 371–382 (1991).

36. G. Stuart, B. Sakmann, Amplification of EPSPs by axosomatic sodium channels in neocortical pyramidal neurons. Neuron 15, 1065–1076 (1995).

37. E. L. Keller, Participation of medial pontine reticular formation in eye movement generation in monkey. J Neurophysiol 37, 316–332 (1974).

38. S. Everling, M. Paré, M. C. Dorris, D. P. Munoz, Comparison of the discharge characteristics of brain stem omnipause neurons and superior colliculus fixation neurons in monkey: implications for control of fixation and saccade behavior. J Neurophysiol 79, 511–528 (1998).

39. C. K. Hauser, D. Zhu, T. R. Stanford, E. Salinas, Motor selection dynamics in FEF explain the reaction time variance of saccades to single targets. Elife 7, (2018).

40. U. K. Jagadisan, N. J. Gandhi, Disruption of Fixation Reveals Latent Sensorimotor Processes in the Superior Colliculus. J Neurosci 36, 6129–6140 (2016).

41. D. P. Hanes, R. H. Wurtz, Interaction of the frontal eye field and superior colliculus for saccade generation. J Neurophysiol 85, 804–815 (2001).

42. T. R. Peel, S. Dash, S. G. Lomber, B. D. Corneil, Frontal eye field inactivation alters the readout of superior colliculus activity for saccade generation in a task-dependent manner. J Comput Neurosci, (2020).

43. B. D. Corneil, E. Olivier, D. P. Munoz, Visual responses on neck muscles reveal selective gating that prevents express saccades. Neuron 42, 831–841 (2004).

44. C. Gu, D. K. Wood, P. L. Gribble, B. D. Corneil, A Trial-by-Trial Window into Sensorimotor Transformations in the Human Motor Periphery. J Neurosci 36, 8273–8282 (2016).

45. A. P. Georgopoulos, A. B. Schwartz, R. E. Kettner, Neuronal population coding of movement direction. Science 233, 1416–1419 (1986).

46. C. Lee, W. H. Rohrer, D. L. Sparks, Population coding of saccadic eye movements by neurons in the superior colliculus. Nature 332, 357–360 (1988).

47. G. Rizzolatti, L. Riggio, I. Dascola, C. Umilta, Reorienting attention across the horizontal and vertical meridians: evidence in favor of a premotor theory of attention. Neuropsychologia 25, 31–40(1987).

48. G. Vigneswaran, R. Philipp, R. N. Lemon, A. Kraskov, M1 corticospinal mirror neurons and their role in movement suppression during action observation. Curr Biol 23, 236–243 (2013).

49. A. Y. Tan, Y. Chen, B. Scholl, E. Seidemann, N. J. Priebe, Sensory stimulation shifts visual cortex from synchronous to asynchronous states. Nature 509, 226–229 (2014).

50. C. Hauptmann, P. A. Tass, Restoration of segregated, physiological neuronal connectivity by desynchronizing stimulation. Journal of neural engineering 7, 056008 (2010).

51. O. V. Popovych, P. A. Tass, Control of abnormal synchronization in neurological disorders. Front Neurol 5, 268 (2014).

52. W. Softky, Sub-millisecond coincidence detection in active dendritic trees. Neuroscience 58, 13–41 (1994).

53. T. R. Stanford, E. G. Freedman, D. L. Sparks, Site and parameters of microstimulation: evidence for independent effects on the properties of saccades evoked from the primate superior colliculus. J Neurophysiol 76, 3360–3381 (1996).

54. H. A. Katnani, N. J. Gandhi, The relative impact of microstimulation parameters on movement generation. J Neurophysiol 108, 528–538 (2012).

55. C. L. Bryant, N. J. Gandhi, Real-time data acquisition and control system for the measurement of motor and neural data. J Neurosci Methods 142, 193–200 (2005).

56. D. L. Sparks, R. Holland, B. L. Guthrie, Size and distribution of movement fields in the monkey superior colliculus. Brain Res 113, 21–34 (1976).

57. V. Mante, D. Sussillo, K. V. Shenoy, W. T. Newsome, Context-dependent computation by recurrent dynamics in prefrontal cortex. Nature 503, 78–84 (2013).

58. M. Rigotti et al., The importance of mixed selectivity in complex cognitive tasks. Nature 497, 585–590(2013).

59. F. P. Ottes, J. A. Van Gisbergen, J. J. Eggermont, Visuomotor fields of the superior colliculus: a quantitative model. Vision Res 26, 857–873 (1986).

60. R. A. Fisher, The use of multiple measurements in taxonomic problems. Ann Eugenic 7, 179–188 (1936).

61. B. W. Matthews, Comparison of the predicted and observed secondary structure ofT4 phage lysozyme. Biochim Biophys Acta 405, 442–451 (1975).

62. S. Boughorbel, F. Jarray, M. El-Anbari, Optimal classifier for imbalanced data using Matthews Correlation Coefficient metric. PLoS One 12, e0177678 (2017).

63. A. Rauch, G. La Camera, H. R. Luscher, W. Senn, S. Fusi, Neocortical pyramidal cells respond as integrate-and-fire neurons to in vivo-like input currents. J Neurophysiol 90, 1598–1612 (2003).

64. M. Usher, J. L. McClelland, The time course of perceptual choice: the leaky, competing accumulator model. Psychol Rev 108, 550–592 (2001).

65. B. A. Purcell et al., Neurally constrained modeling of perceptual decision making. Psychol Rev 117, 1113–1143 (2010).

66. K. F. Wong, A. C. Huk, M. N. Shadlen, X. J. Wang, Neural circuit dynamics underlying accumulation of time-varying evidence during perceptual decision making. Front Comput Neurosci 1, 6 (2007).

67. T. Tsujimoto, A. Jeromin, N. Saitoh, J. C. Roder, T. Takahashi, Neuronal calcium sensor 1 and activity-dependent facilitation of P/Q-type calcium currents at presynaptic nerve terminals. Science 295, 2276–2279 (2002).

68. J. G. Borst, B. Sakmann, Facilitation of presynaptic calcium currents in the rat brainstem. The Journal of physiology 513 (Pt 1), 149–155 (1998).

69. G. G. Turrigiano, K. R. Leslie, N. S. Desai, L. C. Rutherford, S. B. Nelson, Activity-dependent scaling of quantal amplitude in neocortical neurons. Nature 391, 892–896 (1998).

